# A non-catalytic role for RFC in PCNA-mediated processive DNA synthesis

**DOI:** 10.1101/2025.08.08.669392

**Authors:** Gabriella N. L. Chua, Emily C. Beckwitt, Victoria Miller-Browne, Olga Yurieva, Dan Zhang, Bryce J. Katch, John W. Watters, Kaitlin Abrantes, Ryogo Funabiki, Xiaolan Zhao, Michael E. O’Donnell, Shixin Liu

**Affiliations:** Laboratory of Nanoscale Biophysics and Biochemistry, The Rockefeller University, New York, NY, USA; Laboratory of DNA Replication, The Rockefeller University, New York, NY, USA; Howard Hughes Medical Institute, The Rockefeller University, New York, NY, USA; Molecular Biology Program, Memorial Sloan Kettering Cancer Center, New York, NY, USA; Programs in Biochemistry, Cell, and Molecular Biology, Weill Cornell Graduate School of Medical Sciences, New York, NY, USA

**Keywords:** DNA replication, DNA damage, Sliding clamp, Clamp loader, Okazaki fragment, Fill-in synthesis, Genome maintenance, PCNA, RFC, FEN1, Polδ

## Abstract

The ring-shaped sliding clamp PCNA enables DNA polymerases to perform processive DNA synthesis during replication and repair. The loading of PCNA onto DNA is catalyzed by the ATPase clamp loader RFC. Using a single-molecule platform to visualize the dynamic interplay between PCNA and RFC on DNA, we unexpectedly discovered that RFC continues to associate with PCNA after loading, contrary to the conventional view. Functionally, this clamp-loader/clamp complex is required for processive DNA synthesis by polymerase δ (Polδ), as the PCNA-Polδ assembly is inherently unstable. This architectural role of RFC is dependent on the BRCT domain of Rfc1, and mutation of its DNA-binding residues causes sensitivity to DNA damage in vivo. We further showed the FEN1 flap endonuclease can also stabilize the PCNA-Polδ interaction and mediate robust synthesis. Overall, our work revealed that, beyond their canonical enzymatic functions, PCNA-binding proteins harbor non-catalytic functions essential for DNA replication and genome maintenance.

## INTRODUCTION

Genome replication across all domains of life requires DNA sliding clamps that couple with DNA polymerases to achieve processive DNA synthesis [1–3]. In eukaryotes, this activity is fulfilled by proliferating cell nuclear antigen (PCNA), a homotrimer that forms a ring-shaped complex encircling double-stranded (ds) DNA [4, 5]. The closed-ring architecture of PCNA creates a topological challenge for its loading onto DNA, and this challenge is overcome by a clamp-loader complex known as replication factor C (RFC), a heteropentameric ATPase [6]. In the prevailing model, RFC utilizes its ATPase activity to open the PCNA ring and deposit it on a primer-template junction with a 3’ recessed end; upon ATP hydrolysis, RFC is then released from DNA while PCNA is poised to interact with a polymerase for DNA synthesis [7]. However, RFC has also been shown to mediate PCNA loading onto nicked DNA substrates [8–11], suggesting the RFC-PCNA interaction is more nuanced than previously thought.

Besides RFC, PCNA also interacts with many other proteins that play diverse functions ranging from DNA damage repair to chromatin assembly and sister chromatid cohesion [12, 13]. Further, the homotrimeric architecture of PCNA theoretically allows it to bind up to three partners simultaneously, leading to the proposal of a “toolbelt” model. In one rendition of this model, PCNA binds DNA polymerase δ (Polδ), flap endonuclease I (FEN1), and DNA ligase I (Lig1), three enzymes involved in Okazaki fragment maturation during lagging strand synthesis. Nevertheless, whether PCNA binds these proteins simultaneously or sequentially is still under debate [14, 15]. In general, how PCNA recruits, exchanges, and coordinates different proteins and their activities remains poorly understood.

In this study, we used single-molecule correlative force and fluorescence microscopy to visualize the dynamic behavior of PCNA and RFC on DNA. This approach led us to the surprising finding that these two complexes frequently remain associated on DNA even after PCNA loading. The sustained association of the PCNA-RFC assembly with the replication machinery is essential for processive DNA synthesis in vitro. Mutating the domain in RFC required for this association results in impaired genome maintenance in vivo. This work thus reveals an unexpected non-catalytic function of RFC, beyond its canonical clamp-loading activity, and therefore extends the possible functionality of clamp loaders.

## RESULTS

### PCNA and RFC form a long-lived complex that diffuses on dsDNA

We purified *Saccharomyces cerevisiae* PCNA and RFC (**Figure S1A**) and fluorescently labeled each of them with LD655 and Cy3 fluorophores, respectively. Through force manipulation of a 19-kilobase (kb) dsDNA with two pre-engineered nicks by optical tweezers, we generated a gapped DNA substrate containing a 3-kb single-stranded (ss) DNA region flanked by 10-kb and 6-kb dsDNA arms (**Figures S1B** and **S1C**). This gapped DNA, containing both 3’ and 5’ recessed DNA ends, was used to visualize the behavior of PCNA via scanning confocal fluorescence microscopy (**Figure 1A**). To mimic the in vivo scenario, we also included an excess of unlabeled RPA, the eukaryotic ssDNA binding protein. We were able to observe clear PCNA signals on the DNA in the presence of RFC and ATP (**Figures 1B** and **S2A**), whereas PCNA binding was completely abolished when either RFC or ATP was omitted (**Figures S2B** and **S2C**). These results are in agreement with the current understanding of the PCNA loading mechanism and thus validate our experimental platform.

**Figure 1.**
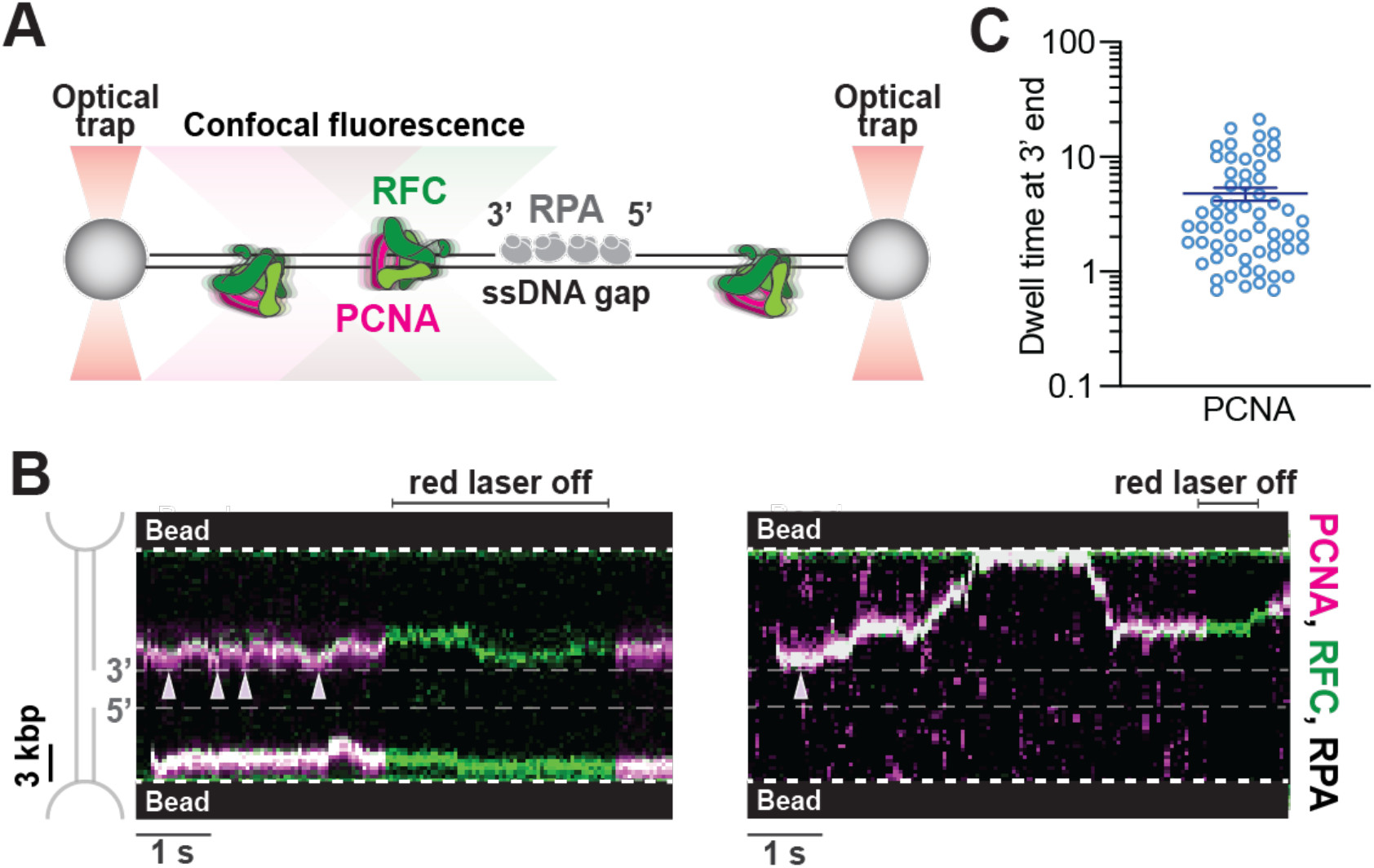
Clamp-loader/clamp complexes bind and slide on dsDNA. (**A**)Schematic of the experimental setup. A single DNA molecule containing a 3-kb ssDNA gap flanked by 10-kb and 6-kb dsDNA arms was tethered between a pair of optically trapped beads through biotin-streptavidin linkage. The tether was moved to a channel containing LD655-PCNA, Cy3-RFC, RPA, and ATP, and their behavior was imaged via dual-color scanning confocal fluorescence microscopy. (**B**)Two representative kymographs showing gapped DNA tethers incubated with 8 nM LD655-PCNA, 8 nM Cy3-RFC, and 100 nM RPA. The red laser was turned off briefly to confirm the presence of the RFC fluorescence signal. Arrows denote the events when PCNA is engaged with the 3’ recessed end of the ssDNA gap. (**C**)Dwell times of PCNA binding events at the 3’ recessed end of the ssDNA gap. Bars represent mean and SEM (*N* = 61 events). See also **Figures S1** and **S2**.

Surprisingly however, our single-molecule data revealed a more complex picture of PCNA behavior on DNA than the prevailing model: rather than stably residing at 3’ recessed DNA ends, PCNA was observed to transiently engage with the 3’ end (white arrows in **Figure 1B**), yielding an average dwell time of 5.4 ± 0.9 s (mean ± SEM) (**Figure 1C**). Most of the time, PCNA stayed on either of the flanking dsDNA arms and displayed a one-dimensional sliding behavior (**Figures 1B** and **S2A**). No PCNA signal was observed within the ssDNA gap region, indicating that RPA-coated ssDNA serves as an efficient barrier against PCNA binding and sliding.

The other striking and unexpected observation from our data was that PCNA and RFC frequently remained associated on DNA and slid together as a complex, evidenced by diffusive trajectories containing both Cy3-RFC and LD655-PCNA signals (**Figures 1B** and **S2A**). This observation stands in contrast to the widely held notion that RFC is immediately released from DNA upon PCNA loading. We hereafter refer to these long-lived RFC-PCNA assemblies as Clamp-Loader/Clamp (CLC) complexes.

### CLCs can be topologically bound to DNA

With the gapped DNA substrate, we frequently observed that the dual-color CLC signal first appeared within a dsDNA region (e.g. **Figures 1B** and **S2A**) rather than at a recessed 3’ or 5’ DNA end. To test whether CLC binding to DNA relies on recessed DNA ends at all, we performed single-molecule experiments using DNA tethers derived from the 48.5-kb-long bacteriophage λ genomic dsDNA, which lacks ssDNA gaps. We observed that CLC complexes can readily bind and slide on this substrate in an RFC- and ATP-dependent manner (**Figures 2A, S2D**, and **S2E**), a fraction of which presumably enter via nicks that may form during purification of the long duplex DNA. Thus, PCNA does not require a recessed end for DNA binding, consistent with previous biochemical data that RFC can load PCNA at a nicked site [8–10]. Of note, we found that RFC alone can also slide on dsDNA in the presence of ATP (**Figure S2F**).

**Figure 2.**
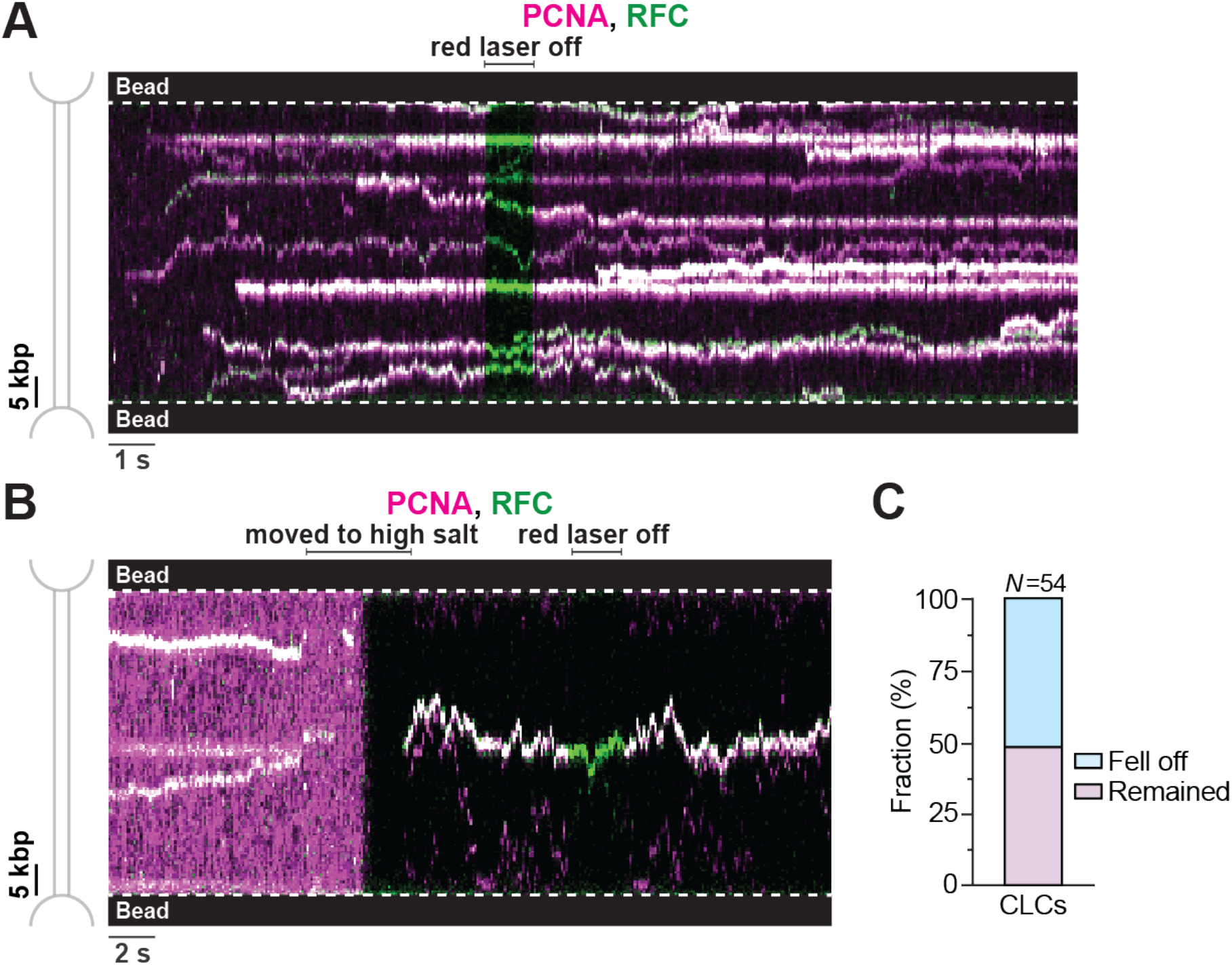
Clamp-loader/clamp complexes can be topologically bound to DNA. (**A**)Representative kymograph showing a double-stranded 11 DNA tether incubated with 8 nM LD655-PCNA, 8 nM Cy3-RFC, and 100 nM RPA with ATP. Red laser was turned off briefly to confirm the presence of green RFC signals. (**B**) Kymograph showing a ds 11 DNA tether first incubated in a channel containing 8 nM LD655-PCNA, 8 nM Cy3-RFC, and 100 nM RPA with ATP and then moved into a separate channel containing a high-salt buffer (500 mM NaCl) with ATP. (**C**) Fraction of clamp-loader/clamp complexes (CLCs) that remained bound to or fell off dsDNA when challenged with high salt (*N* = 54 from 17 independent tethers). See also **Figure S2**.

Next, we asked whether these long-lived CLC complexes contain a topologically closed or open PCNA ring. To this end, we challenged CLC complexes bound to the λ DNA with 500 mM NaCl and found that a substantial fraction of them (~50%) persisted on DNA and continued to slide in this high-salt buffer (**Figures 2B** and **2C**). We presume that this high-salt-resistant population of CLC complexes likely corresponds to those containing a topologically closed PCNA ring encircling the dsDNA.

### CLC formation requires the BRCT domain of RFC

The BRCA1 C-terminal homology (BRCT) domain on the Rfc1 subunit, a region flexibly attached to RFC’s main AAA+ domain core, makes extensive contacts with dsDNA (**Figure 3A**). Therefore, we considered whether there might be persistent association of the CLC with DNA after PCNA loading. To test this, we purified and fluorescently labeled an RFC complex lacking the BRCT domain (RFC^ΔBRCT^) (**Figure S1A**) and examined its behavior with PCNA on the gapped DNA substrate. We found PCNA can be efficiently recruited to DNA by RFC^ΔBRCT^ and then undergo diffusive movements on either 5’ or 3’ dsDNA arms of the ssDNA gap (**Figure 3B**). However, in contrast to the full-length RFC (**Figure 1B**), RFC^ΔBRCT^ was much less frequently observed to colocalize with PCNA (**Figure 3C**), suggesting its eviction from DNA after loading PCNA. The lack of a RFC^ΔBRCT^ fluorescence signal was not due to a low labeling efficiency, as LD555-RFC^ΔBRCT^ can be readily detected (often without colocalized PCNA) within the RPA-coated ssDNA region (**Figure S3**). Moreover, PCNA loaded by RFC^ΔBRCT^ exhibited a much greater diffusion coefficient than those loaded by full-length RFC (**Figure 3D**), which may be explained by the fact that complexes with a larger molecular weight (i.e. CLC compared to PCNA alone) would diffuse more slowly. Together, these results suggest that Rfc1’s BRCT domain is critical for the stable formation of CLC on DNA.

**Figure 3.**
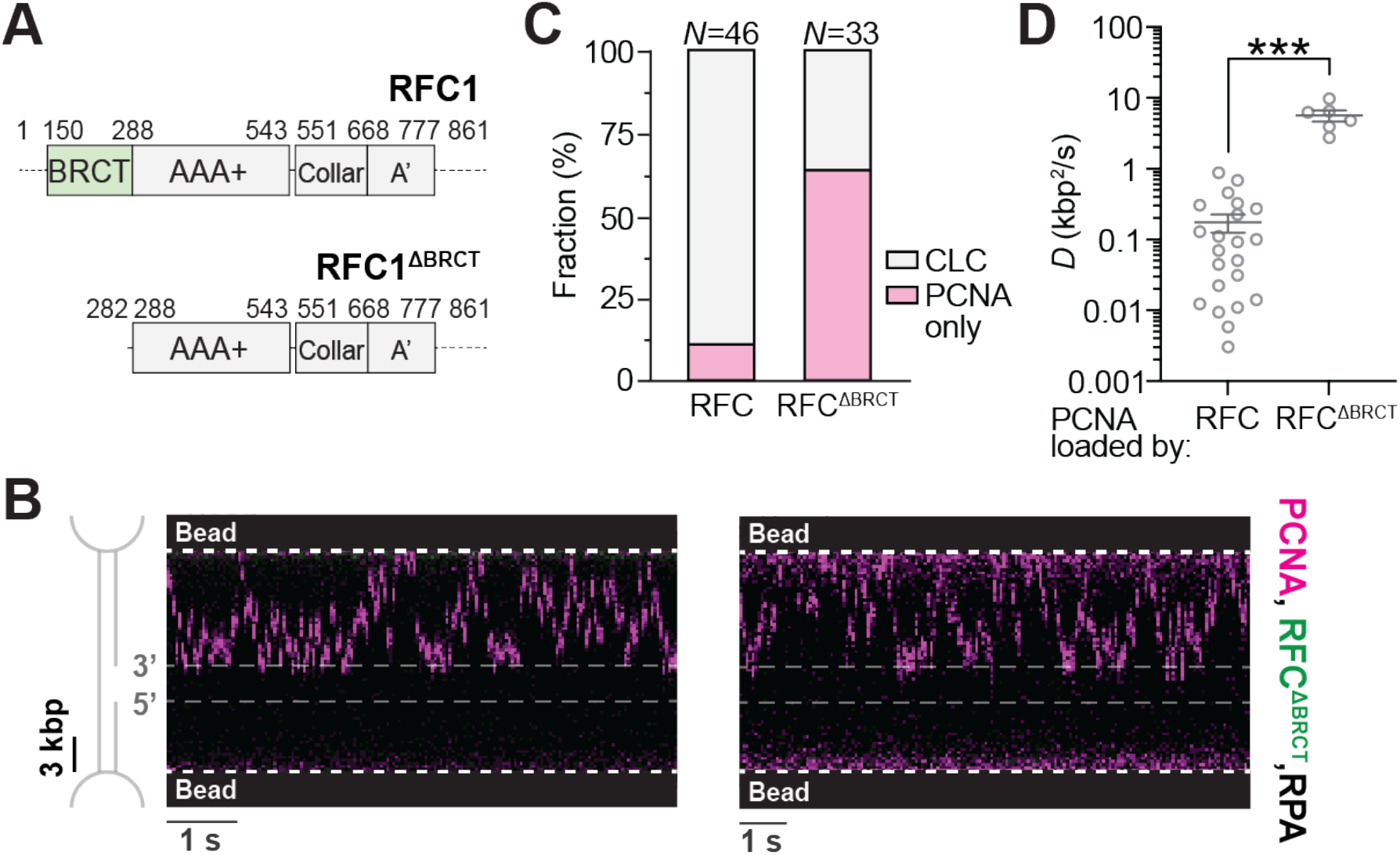
The BRCT domain of RFC promotes CLC formation on DNA. (**A**) Domain structure of Rfc1 and Rfc1^ΔBRCT^. The BRCT domain is highlighted in green. (**B**) Two representative kymographs showing a gapped DNA tether incubated with 8 nM LD655-PCNA, 8 nM LD555-RFC^ΔBRCT^, and 100 nM RPA. (**C**) Fraction of trajectories on gapped DNA containing CLCs or PCNA only when RFC (*N* = 46 from 6 independent tethers) or RFC^ΔBRCT^ (*N* = 33 from 8 independent tethers) was used. (**D**) Diffusion coefficients (*D*) for PCNA trajectories sliding on DNA when RFC (*N* = 22 from 5 independent tethers) or RFC^ΔBRCT^ (*N* = 6 from 6 independent tethers) was used. Bars represent mean and SEM. Significance was calculated using a two-tailed unpaired t-test with Welch’s correction. See also **Figures S1** and **S3**.

### CLC assembles with Polδ to facilitate DNA synthesis

We next explored the functional relevance of the CLC complex during DNA synthesis. To this end, we included Polδ, the DNA polymerase mainly responsible for lagging strand synthesis, in our single-molecule assay with the gapped DNA substrate to observe DNA fill-in synthesis across the ssDNA gap in real time. We first performed this assay using AlexaFluor488-labeled RPA in a distance-clamp mode where the positions of the two traps were fixed. In the presence of PCNA, RFC, Polδ (all unlabeled), and dNTPs, we observed progressive clearance of the RPA signal from the 3’ to the 5’ end of the gap (**Figure 4A**), confirming that the proteins used were active. To visualize the behaviors of PCNA and RFC during the fill-in synthesis, we used unlabeled RPA and instead followed the reaction from the force readings (**Figure 4B**). The conversion from ssDNA to dsDNA causes tether contraction at a starting force above 6 pN and thus a concomitant increase in the DNA tension [16]. We monitored the fluorescence signals from both PCNA and RFC and found that they frequently remained associated and traveled continuously across the gap as the fill-in synthesis progressed (**Figure 4B**). To circumvent complications associated with the varying forces when interpreting the data, we also conducted the experiments in a force-clamp mode where one of the trap positions was adjusted throughout the reaction to maintain a constant DNA tension. Changes in the trap position were converted to the number of nucleotides filled in to track the progression of DNA synthesis (**Figure 4C**). Importantly, RFC was observed along the path of fill-in together with PCNA in the majority of reactions (9 out of 14 tethers), suggesting that CLC frequently assembles with Polδ during DNA synthesis.

**Figure 4.**
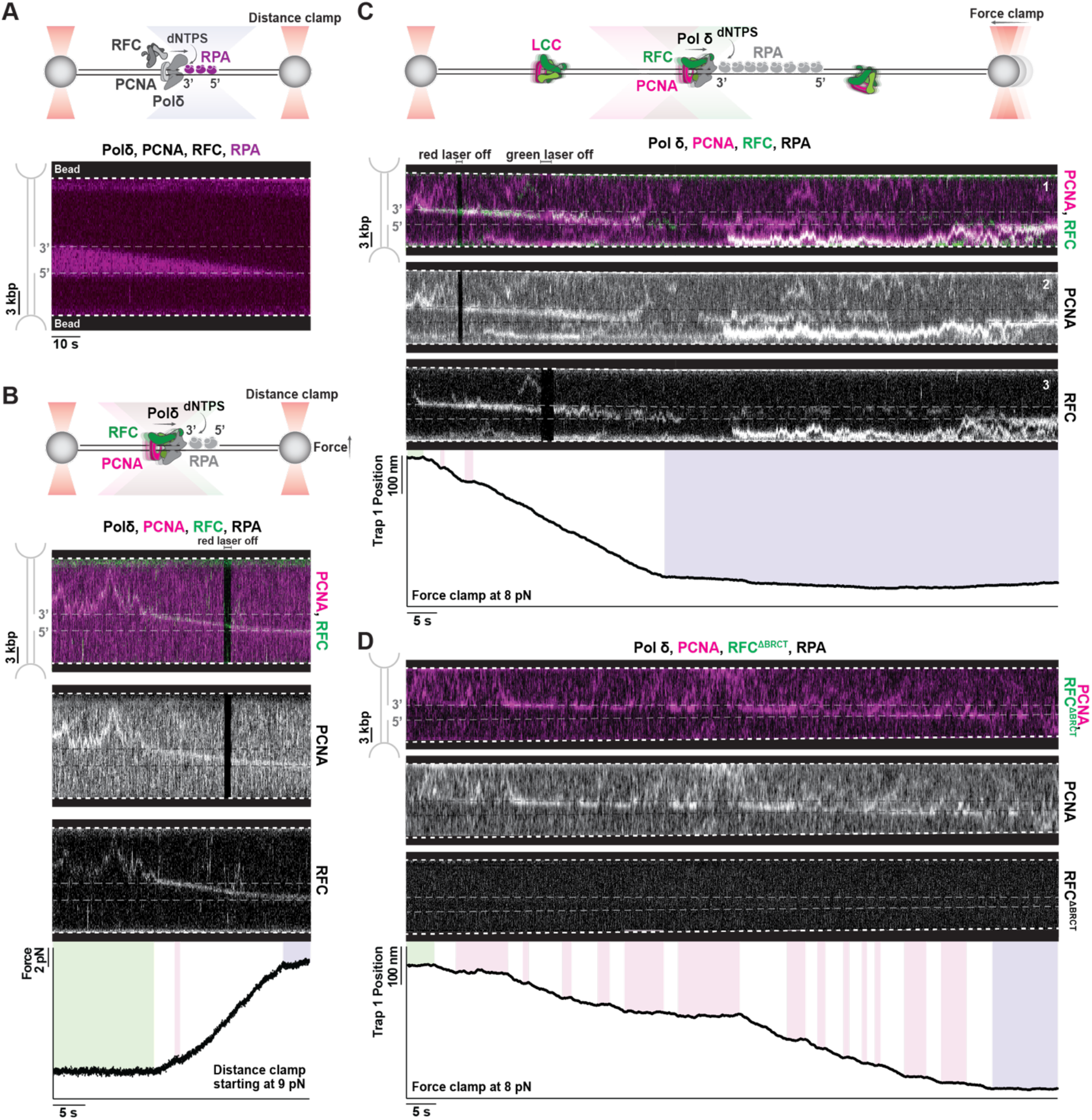
Clamp-loader/clamp complexes assemble with Polδto support DNA synthesis. (**A**)(Top) Schematic of experimental setup. A single gapped DNA molecule was tethered between a pair of optically trapped beads held at a constant distance. The tether was moved to a channel containing Polδ, PCNA, RFC, and AF488-RPA with ATP and dNTPs. The behavior of RPA was imaged using scanning confocal fluorescence microscopy. (Bottom) Kymograph showing RPA clearing during fill-in of the ssDNA gap when incubated with 20 nM Polδ, 5 nM PCNA, 5 nM RFC, and 20 nM AF488-RPA. (**B**) (Top) Schematic of experimental setup where the distance between beads was held constant starting at a force of 9 pN, and fill-in was monitored by an increase in the force as the tether shortened. (Bottom) Kymograph and associated force changes showing fill-in of a gapped DNA tether incubated with 20 nM Polδ, 5 nM LD655-PCNA, 5 nM Cy3-RFC, and 100 nM RPA. Red laser was turned off briefly, and PCNA and RFC fluorescence channels are also separately shown to confirm the presence of each signal. (**C**) (Top) Schematic of experimental setup where the force was held constant, and fill-in was monitored by the movement of one of the traps as the tether shortened. (Bottom) Kymograph and the associated trap position changes at a constant force of 8 pN showing fill-in of a gapped DNA tether incubated in the same conditions as (**B**). (**D**)Kymograph and associated trap position changes at a constant force of 8 pN showing fill-in of a gapped DNA tether incubated with 20 nM Polδ, 5 nM LD655-PCNA, 5 nM LD555-RFC^ΔBRCT^, and 100 nM RPA. In (**B-D**), pink boxes denote periods of inactive synthesis, green box denotes prior to start of fill-in, and purple box denotes completion of fill-in. See also **Figures S4** and **S5**.

The finding that RFC remains part of the replication machinery begs the question of whether it can contribute to DNA synthesis. To answer this question, we took advantage of the RFC^ΔBRCT^ variant, which we showed above to be proficient in loading PCNA onto DNA but deficient in forming CLC complexes. We performed the single-molecule fill-in assay with RFC^ΔBRCT^. In contrast to full-length RFC, RFC^ΔBRCT^ was never observed to co-travel with PCNA along the DNA fill-in path (0 out of 20 tethers). PCNA alone was still observed to translocate across the ssDNA gap, but frequently slid back onto the dsDNA (**Figures 4D** and **S4A**). Strikingly, whenever PCNA left the path of synthesis (i.e. 3’ end of the filled gap), we observed a concomitant pause in the synthesis activity (pink boxes in **Figures 4D** and **S4A**), confirming that PCNA presence at a 3’ terminus is required for processive Polδ activity. We quantified the fill-in synthesis activity in the presence of RFC^ΔBRCT^ or full-length RFC. The fraction of time that the fill-in trajectories spent in an inactive period was significantly higher for RFC^ΔBRCT^ than for full-length RFC (**Figure 5A**). Moreover, a substantial fraction of fill-in trajectories did not complete the entire 3-kb ssDNA track within a maximum allotted time window (350 s) n when PCNA was loaded by RFC^ΔBRCT^, whereas virtually all trajectories completed the reaction when full-length RFC was used (**Figure 5B**).

**Figure 5.**
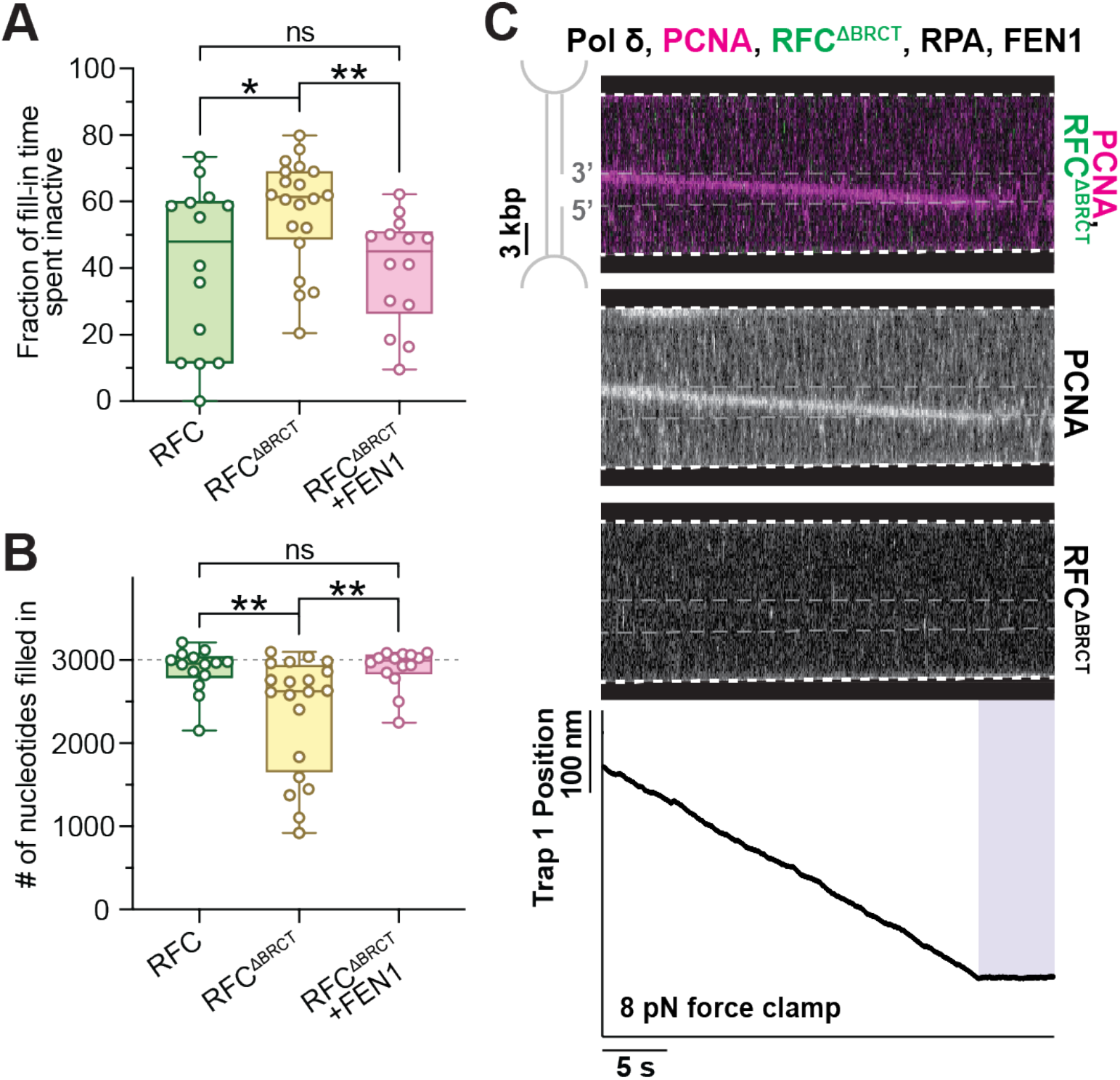
FEN1 rescues deficient fill-in associated with unstable Polδ-PCNA. (**A**) Amount of time spent in an inactive state for reactions incubated with RFC (*N* = 27 from 15 independent tethers), RFC^ΔBRCT^ (*N* = 48 from 20 independent tethers), or RFC^ΔBRCT^ with FEN1 (*N* = 22 from 18 independent tethers). (**B**) Number of nucleotides filled in per tether for reactions incubated with RFC (*N* = 15 from 15 independent tethers), RFC^ΔBRCT^ (*N* = 20 from 20 independent tethers), and RFC^ΔBRCT^ with FEN1 (*N* = 18 from 18 independent tethers). In (**A**) and (**B**), box boundaries represent 25^th^ to 75^th^ percentiles, middle bar represents median, and whiskers represent minimum and maximum values. Significance was calculated using a two-tailed unpaired *t*-test with Welch’s correction. (**C**)Kymograph and associated trap position changes at constant force of 8 pN showing fill-in of a gapped DNA tether incubated with 20 nM Polδ, 5 nM LD655-PCNA, 5 nM LD555-RFC^ΔBRCT^, 15 nM FEN1, and 100 nM RPA. PCNA and RFC^ΔBRCT^ fluorescence channels are separately shown. Purple box denotes completion of fill-in. See also **Figures S5** and **S6**.

To assess whether Polδ remains at the 3’ terminus when PCNA has left, we conducted experiments where we first allowed the fill-in reaction to start in a channel containing Polδ, RFC^ΔBRCT^ and PCNA; once we observed a pause in the synthesis activity, we moved the tether to a different channel lacking free Polδ in solution (**Figure S4B**). The fill-in reaction was never observed to resume (0 out of 7 tethers) even though PCNA repeatedly engaged with the 3’ end, indicating that the original Polδ has also dissociated, and a new polymerase needs to be recruited to continue the reaction. Together, these results demonstrate that besides its catalytic function in loading PCNA onto DNA, RFC also fulfills an architectural function in stabilizing the PCNA-Polδ interaction—which is lost in the RFC^ΔBRCT^ variant—thereby enhancing the DNA synthesis efficiency.

### FEN1 and RFC share a non-catalytic role in DNA synthesis

The differential ability of RFC and RFC^ΔBRCT^ to support processive fill-in synthesis points to the inherent instability of the PCNA-Polδ complex at the growing 3’ end. Given the many binding partners of PCNA [13], we surmised that other factors may also fulfill the same function as RFC in stabilizing PCNA-Polδ interaction. The flap-processing endonuclease FEN1 has been shown to form a ternary complex with PCNA and Polδ in the human system [17]. We therefore sought to examine whether FEN1 can fulfill a similar function as RFC in stabilizing PCNA-Polδ interaction and promoting processive synthesis. We performed the single-molecule fill-in experiments in the presence of FEN1 and RFC^ΔBRCT^, the latter of which is required for PCNA loading but cannot sustain frequent, stable CLC formation on DNA. Remarkably, we found that FEN1 largely rescued the deficiency of RFC^ΔBRCT^ in supporting processive synthesis, increasing the completion rate and decreasing the fraction of inactive periods to levels comparable to those generated in the presence of full-length RFC (**Figures 5A** and **5B**). Indeed, when FEN1 was present, PCNA remained on the fill-in path and rarely slid back onto the dsDNA (**Figure 5C**).

These observations suggest that, even though RFC and FEN1 each have their own catalytic functions in lagging strand synthesis, they also share a non-catalytic role in stabilizing the PCNA-Polδ assembly. To gain further insight into this redundant structural function, we used AlphaFold3 to model the architecture of the PCNA-RFC-Polδ complex with a 3’ recessed DNA (**Figure S5A**). The predicted structure showed PCNA and Polδ in similar positions on the DNA as in a cryoelectron microscopy (cryo-EM) structure of the binary complex poised for synthesis [18]. It further showed that RFC occupies the space between the polymerase and one protomer of PCNA. Interestingly, FEN1 was observed to occupy the same space in the cryo-EM structure of the PCNA-FEN1-Polδ complex (**Figure S5B**) [17], reinforcing the notion of FEN1 and RFC adopting redundant structural roles in stabilizing the fill-in machinery.

Next, we asked whether FEN1 and RFC compete for PCNA binding. We first used mass photometry to measure the distribution of complex sizes with different combinations of proteins. When PCNA was incubated with RFC alone, a predominant mass peak corresponding to the CLC complex was observed (**Figure S6A**). However, when an excess of FEN1 was added, the CLC peak diminished and complexes corresponding to PCNA bound to one or more copies of FEN1 were observed (**Figure S6B**). We further performed single-molecule experiments to evaluate CLC formation on DNA in the presence of FEN1. We found that the likelihood of PCNA co-sliding with RFC on dsDNA was drastically reduced by the addition of excess FEN1 (**Figures S6C** and **S6D**). These results indicate that excess FEN1 promotes RFC eviction from PCNA both on and off DNA.

### DNA-binding residues in the BRCT domain are essential for genome maintenance

Our single-molecule data uncovered an underappreciated structural role of RFC in supporting DNA synthesis via the formation of CLC complexes and the importance of Rfc1’s BRCT domain in this activity. To corroborate these findings, we examined the effects of BRCT mutants on genome maintenance in vivo. We generated *S. cerevisiae* cells wherein the endogenous *RFC1* gene was replaced with a mutant allele in which the sequence encoding the BRCT domain was deleted (*rfc1-BRCTΔ*). We also generated cells containing a mutant *rfc1* allele in which 11 residues in the BRCT domain that have been identified to contact dsDNA [8–10] were mutated into alanine (*rfc1-11A*). The wild-type Rfc1, Rfc1-BRCTΔ, and Rfc1-11A proteins were tagged with a TAP tag at their C-termini to assess protein levels. Neither mutant protein showed a significantly different expression level compared to the wild-type (**Figure S7A**). When cells were challenged with different DNA damaging agents, *rfc1-BRCTΔ* cells showed sensitivity to the DNA-methylating agent methyl methanesulfonate (MMS), but not to hydroxyurea (HU) that blocks dNTP synthesis or to camptothecin (CPT) that inhibits the DNA topoisomerase 1 (**Figure 6A**) [19]. The unique sensitivity to MMS is consistent with previous studies using plasmid-born *rfc1* mutants with the BRCT domain or a larger N-terminal region deleted [10, 20]. *rfc1-11A* showed even more striking sensitivity to MMS (**Figure 6A**). As a control, wild-type Rfc1 tagged with TAP did not show sensitivity to MMS (**Figures 6A** and **S7B**), suggesting that the mutants’ defects were not caused by tagging. The sensitivity profile of *rfc1-BRCTΔ* and *rfc1-11A* resembles those caused by deficiencies in proteins involved in lagging strand synthesis or base excision repair [21–23], providing evidence that the BRCT domain and its DNA-binding sites contribute to these processes. The less pronounced defect caused by *rfc1-BRCTΔ* compared with *rfc1-11A* can be related to higher protein level of the former mutant (**Figure S7A**), though other explanations are possible.

**Figure 6.**
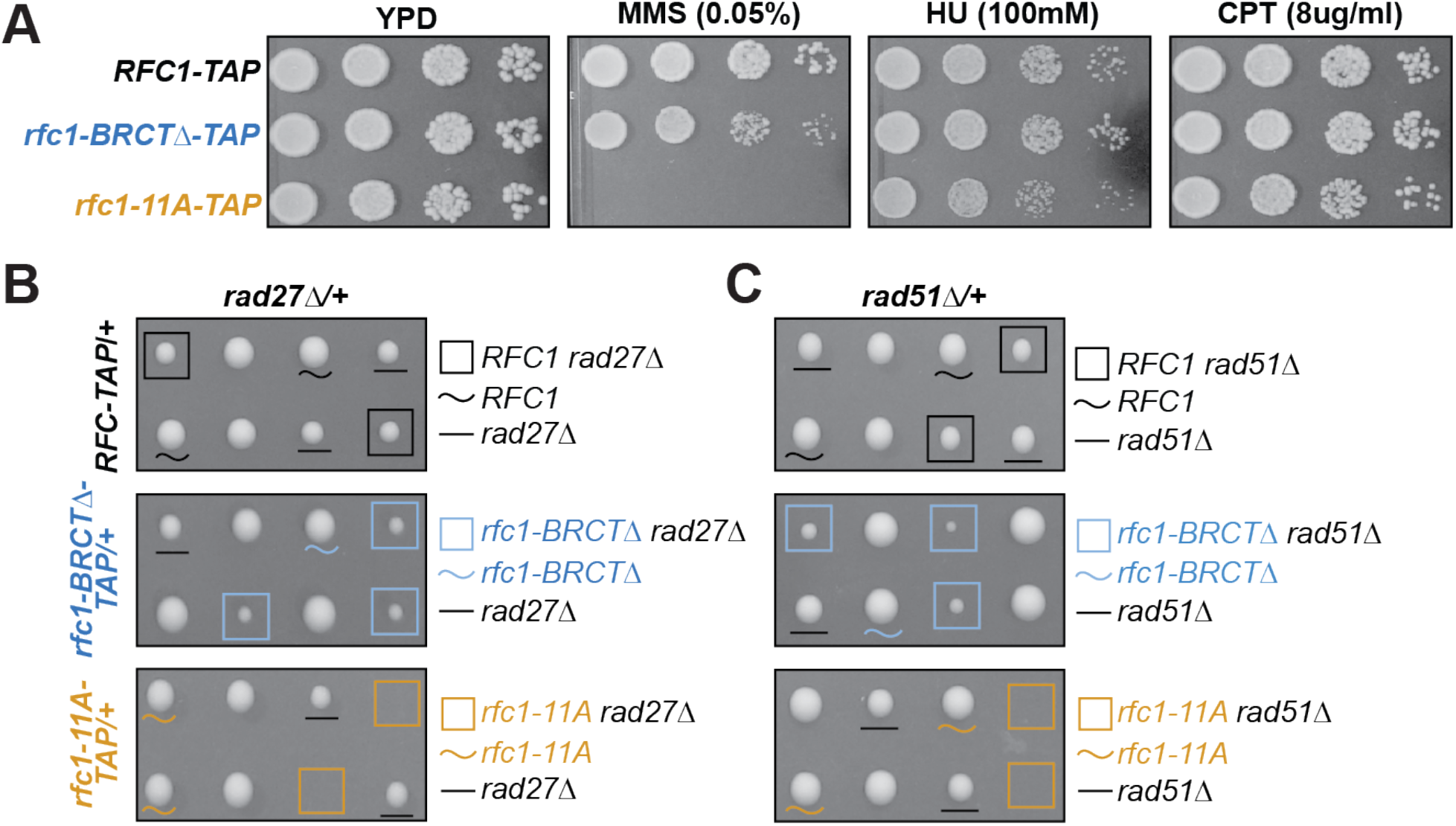
Mutating Rfc1’s BRCT domain or its DNA-binding residues sensitizes cells to genotoxins and the lack of FEN1 or Rad51. (**A**) Cells harboring *rfc1-BRCTΔ* or *rfc1-11A* were grown in the absence and presence of MMS, HU, or CPT. Ten-fold serial dilution of cells were used. (**B**) Genetic interactions of *rfc1-BRCTΔ* and *rfc1-11A* with FEN1 (Rad27) deletion. Diploid cells with indicated genotypes were dissected and two representative tetrads, each containing 4 spore clones for each diploid are shown. Symbols denote spore clones with the indicated genotypes, and wild-type spore clones are not marked. Size comparison among spore clones was conducted within the same plate. (**C**) Genetic interactions of *rfc1-BRCTΔ* and *rfc1-11A* with Rad51 deletion. Experiments were done and data are presented as in (**B**). See also **Figure S7** and **Table S1**.

### FEN1 deletion is synthetically lethal in combination with RFC BRCT DNA-binding mutations

To further explore the biological pathways that the Rfc1 BRCT domain can affect, we examined the genetic interactions of *rfc1-BRCTΔ* and *rfc1-11A* with a panel of deletion mutants that impact different genome maintenance pathways. Considering FEN1 could rescue deficient fill-in caused by RFC^ΔBRCT^ in our single-molecule experiments, we first deleted the *RAD27* gene that encodes FEN1 in yeast and performed tetrad analyses of diploid cells heterozygous for *rfc1-BRCTΔ* or *rfc1-11A* and for *rad27Δ*. We found both *rfc1* mutants exhibited negative interactions with *rad27Δ*. While *rfc1-BRCTΔ* reduced the sizes of the *rad27Δ* spore clones, *rfc1-11A* was synthetically lethal with *rad27Δ* (**Figure 6B**). As a control, the TAP-tagged wild-type Rfc1 did not affect the growth of *rad27Δ* spore clones (**Figure 6B**, top). In conjunction with our single-molecule results, the simplest interpretation of these negative genetic interactions is that the Rfc1 BRCT domain and FEN1 both affect lagging strand DNA synthesis.

Next, we examined the genetic interaction between BRCT mutations and deletion of the gene encoding for the recombinase Rad51, which can mediate gap repair when lagging strand synthesis is impaired [24]. Similar to the *rad27Δ* results, both *rfc1-BRCTΔ* and *rfc1-11A* substantially slowed the growth of *rad51Δ*-containing spore clones, with *rfc1-11A* exhibiting a more dramatic effect (**Figure 6C**). Furthermore, we deleted the gene encoding the DNA damage checkpoint kinase Mec1 (*mec1Δ*) that can be activated by lagging strand synthesis defects [25]. The *SML1* gene was also deleted to support *mec1Δ* cell viability as *mec1Δ* is lethal [26]. We found that *rfc1-11A* showed a strong negative interaction with *mec1Δ sml1Δ* (**Figure S7C**, left). Finally, we deleted genes encoding RNaseH1 and RnaseH2, which remove DNA:RNA hybrids that can form during lagging strand synthesis [27]. We found the double mutant (*rnh1Δ rnh201Δ*), but not single mutants, had a synthetic lethal relationship with *rfc1-11A* (**Figure S7C**, right). The BRCT deletion mutant *rfc1-BRCTΔ* also caused a reduction in the size of spore clones containing *mec1Δ sml1Δ* or *rnh1Δ rnh201Δ*, although to a lesser degree compared to *rfc1-11A* (**Figure S7C**). This growth defect became more evident upon MMS treatment (**Figure S7D**). Collectively, the genetic profiles provide evidence for the importance of the Rfc1 BRCT domain, particularly its DNA binding sites, in genome maintenance related to lagging strand synthesis.

## DISCUSSION

The conventional model depicts that RFC predominantly deposits PCNA at a recessed 3’ DNA end and gets evicted from the DNA soon after. Contrary to this model, through direct real-time visualization, our single-molecule data reveal that RFC-mediated PCNA loading on DNA does not require a recessed end, echoing recent biochemical and structural studies [8–10]. Moreover, we made the surprising finding that RFC tends to remain associated with PCNA after DNA loading, together forming a clamp-loader/clamp (CLC) complex that can be topologically linked to and slide on dsDNA (**Figure 7**). We further found evidence for the functional importance of CLC complexes in vitro and in vivo, specifically in stabilizing the replication machinery during DNA synthesis. As such, by staying engaged with PCNA and the polymerase, RFC fulfills a non-catalytic, architectural role after accomplishing its ATP-dependent clamp-loading function. Notably, it was recently reported that under replication stress when excess Okazaki fragments are produced, free pools of both PCNA and RFC are depleted[28, 29]. This result would be difficult to rationalize if RFC solely acts as a catalytic enzyme; rather depletion of free PCNA and RFC in cells can be explained by our finding that CLCs accumulate on DNA.

**Figure 7.**
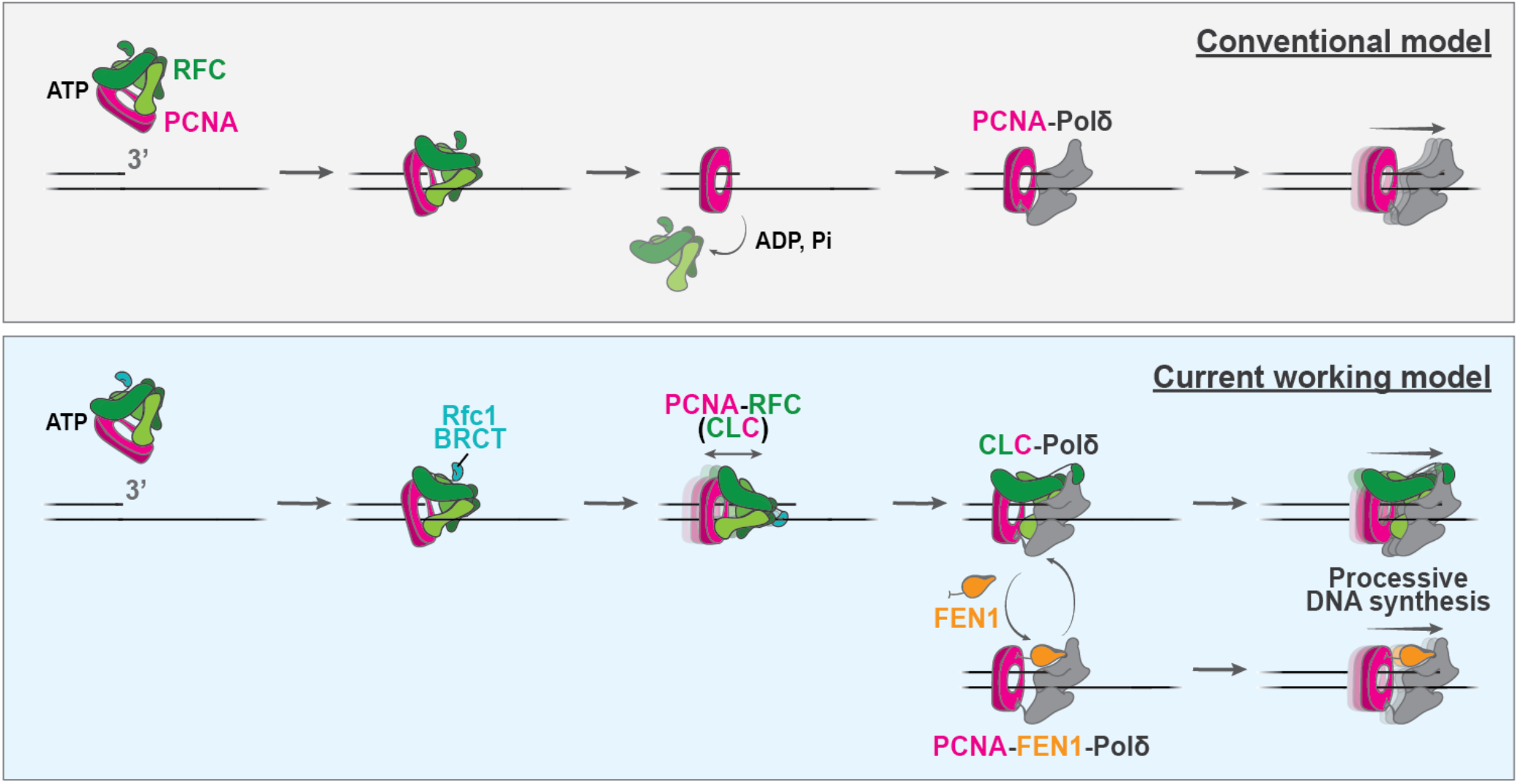
RFC and FEN1 possess non-catalytic roles that enable processive DNA synthesis by Polδ. (**B**) (Top) In the conventional model, RFC binds PCNA in the presence of ATP, loads it onto a primer-template junction with a 3’ recessed end, and releases itself from the DNA upon ATP hydrolysis, leaving PCNA on the DNA to interact with a polymerase for DNA synthesis. (Bottom) Based our findings in this work, we propose a revised model for clamp loading. RFC loads PCNA onto DNA and frequently remains associated, which is dependent on an intact Rfc1 BRCT domain. The clamp-loader/clamp (CLC) complex can slide on duplex DNA until engaging with a polymerase to initiate synthesis. RFC can be replaced by FEN1 for PCNA binding. The PCNA-RFC or PCNA-FEN1 complex can assemble with Polδ, leading to stable PCNA-Polδinteraction and processive fill-in synthesis over long ssDNA tracks.

Our work sheds new light on the role of the Rfc1 BRCT domain in RFC function. It has been shown that BRCT deletion has little effect on the PCNA loading activity of RFC, but results in shorter Okazaki fragment lengths in vitro and elevated sensitivity to certain DNA damage agents such as MMS [9]. Our single-molecule results suggest that these defects are in least in part due to the inability of RFC^ΔBRCT^ to form stable CLC complexes with PCNA, which diminishes the processivity of DNA synthesis. The genotoxic sensitivity profile that we obtained for yeast cells lacking DNA-binding-competent Rfc1 BRCT mimics those for mutants deficient in lagging strand synthesis [10, 21, 22, 30]. Consistent with this observation, we detected negative genetic interactions between the BRCT mutants and mutants of proteins involved in lagging strand synthesis (FEN1, RNaseH1/2) or those that respond to lagging strand synthesis defects (Rad51, Mec1). It will be interesting to assess the contribution of the Rfc1 BRCT and the CLC complex to DNA synthesis performed during other genome maintenance processes, such as the base excision repair pathway wherein PCNA-mediated DNA synthesis is also important [20]. In addition, as the BRCT truncation of the human RFC1 protein was found in patients with Hutchinson-Gilford progeria syndrome [31], the structural role of RFC via its BRCT in DNA synthesis may be implicated in the pathology of this disease.

The single-molecule assays developed here can be readily used to examine other sliding clamp-clamp loader/unloader pairs including Ctf18, Elg1, and Rad24-RFC [32]. Additionally, it will be interesting to examine whether the persistent formation of CLC can enhance the processivity of other DNA polymerases, including the leading-strand polymerase Polε and various translesion polymerases, especially where the fill-in of long ssDNA gaps is required [33].

Finally, we showed that RFC is not the only PCNA-binding factor that can stabilize the PCNA-Polδ assembly: FEN1 proficiently rescues the replication defect caused by RFC-BRCT mutation and promotes processive synthesis. Given that the interaction with PCNA is known to stimulate FEN1’s nuclease activity [34], we envision that the replacement of RFC for FEN1 during fill-in synthesis (**Figure 7**) maintains the structural integrity of the replication machinery and poises FEN1 for flap processing, thus coupling its non-catalytic and catalytic functions. It is conceivable that other PCNA-binding partners can serve a similar structural role in addition to their canonical functions in DNA replication, repair, and chromatin maintenance. This work offers an experimental platform to visualize and dissect their dynamic interaction and exchange with PCNA [35].

## ACKNOWLEDGEMENTS

We thank L. Vostal (Rockefeller University) for help with the mass photometry experiments, Doston Karimov, Sofya Ignatyeva, and Jian Zheng in the Zhao lab for help with tetrad dissections and spot assays, and members of the Liu Lab for critical feedback. V.M.-B. acknowledges a training grant from NIGMS awarded to the Molecular Biophysics Program at Weill Cornell Graduate School (T32GM132081). This work was funded by NIH (R01GM149862 to S.L., R35GM148159 to M.E.O., R35GM145260 to X.Z.). S.L. acknowledges support from the Alfred P. Sloan Foundation and the Marlene Hess Center for Research on Women’s Health and Biomedicine. M.E.O. is a Howard Hughes Medical Institute investigator.

## AUTHOR CONTRIBUTIONS

G.N.L.C. conceived the project and performed the single-molecule experiments under the guidance of S.L. and M.E.O.. E.C.B. prepared the DNA substrate and several fluorescently labeled proteins. V. M.-B. generated the mutant yeast stains and performed genetic and immunoblotting assays under the guidance of X.Z.. O.Y. and D.Z. purified the proteins. B.J.K. and J.W.W. wrote the scripts for and assisted with single-molecule data analyses. K.A. and R.F. assisted with single-molecule data collection. G.N.L.C., V. M-B. S.L., M.E.O., and X.Z. wrote the manuscript.

## REFERENCES

1. Stillman, B., Smart machines at the DNA replication fork. Cell, 1994. 78(5): p. 725–728.

2. Benkovic, S.J., A.M. Valentine, and F. Salinas, Replisome-Mediated DNA Replication. Annual Review of Biochemistry, 2001. 70(Volume 70, 2001): p. 181–208.

3. Johnson, A. and M. O’Donnell, Cellular DNA replicases: components and dynamics at the replication fork. Annu Rev Biochem, 2005. 74: p. 283–315.

4. Krishna, T.S.R., et al., Crystal structure of the eukaryotic DNA polymerase processivity factor PCNA. Cell, 1994. 79(7): p. 1233–1243.

5. Gulbis, J.M., et al., Structure of the C-terminal region of p21(WAF1/CIP1) complexed with human PCNA. Cell, 1996. 87(2): p. 297–306.

6. Bowman, G.D., M. O’Donnell, and J. Kuriyan, Structural analysis of a eukaryotic sliding DNA clamp-clamp loader complex. Nature, 2004. 429(6993): p. 724–30.

7. Bell, S.P. and K. Labib, Chromosome Duplication in Saccharomyces cerevisiae. Genetics, 2016. 203(3): p. 1027–67.

8. Zheng, F., et al., Cryo-EM structures reveal that RFC recognizes both the 3′-and 5′-DNA ends to load PCNA onto gaps for DNA repair. eLife, 2022. 11: p. e77469.

9. Schrecker, M., et al., Multistep loading of a DNA sliding clamp onto DNA by replication factor C. eLife, 2022. 11: p. e78253.

10. Liu, X., et al., A second DNA binding site on RFC facilitates clamp loading at gapped or nicked DNA. Elife, 2022. 11.

11. Yao, N.Y., et al., Mechanism of proliferating cell nuclear antigen clamp opening by replication factor C. J Biol Chem, 2006. 281(25): p. 17528–17539.

12. Boehm, E.M., M.S. Gildenberg, and M.T. Washington, Chapter Seven - The Many Roles of PCNA in Eukaryotic DNA Replication, in The Enzymes, L.S. Kaguni and M.T. Oliveira, Editors. 2016, Academic Press. p. 231–254.

13. Moldovan, G.-L., B. Pfander, and S. Jentsch, PCNA, the Maestro of the Replication Fork. Cell, 2007. 129(4): p. 665–679.

14. Beattie, T.R. and S.D. Bell, Coordination of multiple enzyme activities by a single PCNA in archaeal Okazaki fragment maturation. Embo j, 2012. 31(6): p. 1556–67.

15. Dovrat, D., et al., Sequential switching of binding partners on PCNA during in vitro Okazaki fragment maturation. Proc Natl Acad Sci U S A, 2014. 111(39): p. 14118–23.

16. Wasserman, M.R. and S. Liu, A Tour de Force on the Double Helix: Exploiting DNA Mechanics To Study DNA-Based Molecular Machines. Biochemistry, 2019. 58(47): p. 4667–4676.

17. Lancey, C., et al., Structure of the processive human Pol δholoenzyme. Nature Communications, 2020. 11(1): p. 1109.

18. Zheng, F., et al., Structure of eukaryotic DNA polymerase δbound to the PCNA clamp while encircling DNA. Proc Natl Acad Sci U S A, 2020. 117(48): p. 30344–30353.

19. Pommier, Y., et al., Roles of eukaryotic topoisomerases in transcription, replication and genomic stability. Nat Rev Mol Cell Biol, 2016. 17(11): p. 703–721.

20. Gomes, X.V., S.L. Gary, and P.M. Burgers, Overproduction in Escherichia coli and characterization of yeast replication factor C lacking the ligase homology domain. J Biol Chem, 2000. 275(19): p. 14541–9.

21. Gameiro, E., et al., Genome-wide conditional degron libraries for functional genomics. J Cell Biol, 2025. 224(2).

22. Parsons, A.B., et al., Integration of chemical-genetic and genetic interaction data links bioactive compounds to cellular target pathways. Nat Biotechnol, 2004. 22(1): p. 62–9.

23. Oughtred, R., et al., The BioGRID interaction database: 2019 update. Nucleic Acids Res, 2019. 47(D1): p. D529–d541.

24. Branzei, D. and B. Szakal, DNA damage tolerance by recombination: Molecular pathways and DNA structures. DNA Repair (Amst), 2016. 44: p. 68–75.

25. Lanz, M.C., D. Dibitetto, and M.B. Smolka, DNA damage kinase signaling: checkpoint and repair at 30 years. Embo j, 2019. 38(18): p. e101801.

26. Zhao, X., E.G. Muller, and R. Rothstein, A suppressor of two essential checkpoint genes identifies a novel protein that negatively affects dNTP pools. Mol Cell, 1998. 2(3): p. 329–40.

27. Cerritelli, S.M. and R.J. Crouch, Ribonuclease H: the enzymes in eukaryotes. Febs j, 2009. 276(6): p. 1494–505.

28. Bertolin, A.P., et al., The DNA replication checkpoint prevents PCNA/RFC depletion to protect forks from HLTF-induced collapse in human cells. Molecular Cell, 2025. 85(13): p. 2474–2486.e6.

29. Canal, B., et al., The DNA replication checkpoint limits Okazaki fragment accumulation to protect and restart stalled forks. Molecular Cell, 2025. 85(13): p. 2462–2473.e6.

30. Chang, M., et al., A genome-wide screen for methyl methanesulfonate-sensitive mutants reveals genes required for S phase progression in the presence of DNA damage. Proc Natl Acad Sci U S A, 2002. 99(26): p. 16934–9.

31. Tang, H., et al., Replication factor C1, the large subunit of replication factor C, is proteolytically truncated in Hutchinson-Gilford progeria syndrome. Aging Cell, 2012. 11(2): p. 363–5.

32. Li, H., M. O’Donnell, and B. Kelch, Unexpected new insights into DNA clamp loaders: Eukaryotic clamp loaders contain a second DNA site for recessed 5’ ends that facilitates repair and signals DNA damage: Eukaryotic clamp loaders contain a second DNA site for recessed 5’ ends that facilitates repair and signals DNA damage. Bioessays, 2022. 44(11): p. e2200154.

33. Goodman, M.F. and R. Woodgate, Translesion DNA polymerases. Cold Spring Harb Perspect Biol, 2013. 5(10): p. a010363.

34. Gomes, X.V. and P.M. Burgers, Two modes of FEN1 binding to PCNA regulated by DNA. Embo j, 2000. 19(14): p. 3811–21.

35. Choe, K.N. and G.-L. Moldovan, Forging Ahead through Darkness: PCNA, Still the Principal Conductor at the Replication Fork. Molecular Cell, 2017. 65(3): p. 380–392.

