## Supplementary material for "A non-catalytic role for RFC in PCNA-mediated processive DNA synthesis": Methods and Supplemental Figures

#### Protein preparation

##### *PCNA*

*S. cerevisiae* S6-tagged PCNA was expressed and purified using a previously described protocol [1]. Briefly, a 3X FLAG tag, followed by an S6-tag was added to the N-terminus of S.c. PCNA encoded by *Pol30*. The S.c. FLAG-S6-PCNA encoding plasmid (deposited to Addgene # 23471) was transformed into BLR(DE3) cells which were grown to OD 0.6 at 37°C in LB media. Cells were then cooled to 15°C and induced for expression using 1 mM IPTG for 18 h at 15°C before harvesting by centrifugation. Cells were disrupted by French Press using buffer A (20 mM Tris-Cl pH 7.5, 1 mM DTT, 0.1 mM EDTA, 10% glycerol) and the cell lysate was clarified by centrifugation. The cell lysate was applied to a FLAG affinity column equilibrated in buffer A, washed extensively with buffer A + 500 mM NaCl, then eluted using buffer A + 300 mM NaCl and 2 μM FLAG-peptide. Eluted fractions were dialyzed against buffer A + 50 mM NaCl then purified from excess peptide using a Q Sepharose column eluted with a 100-600 mM NaCl gradient in Buffer A. Fractions containing FLAG-S6-PCNA were pooled, flash frozen and stored at -80 °C.

##### *RFC complexes*

*S. cerevisiae* RFC containing full-length Rfc1 having an N-terminal 3XFLAG tag followed by an S6 labeling sequence was cloned into pRS405 (gal) (deposited to Addgene as # 239472) and integrated into a C600 yeast strain [2]. The strain first modified by integration of plasmids encoding the Rfc2-5 subunits under control of the Gal1/10 promotor (i.e. S.c. RFC[2+5}-pRS403/Gal (deposited to Addgene # 239474) and S.c. RFC[3+4]-pRS402/Gal (deposited to Addgene # 239475). A 1 L overnight culture of cells were grown in the appropriate drop out media at 30°C and then inoculated into 24 2L flasks containing 1L YP media and grown overnight at 30°C until the cell culture reached an O.D. = 0.6. Cells were then induced with Gal, allowed to incubate a further 6 h at 30°C, then harvested by centrifugation. Harvested cells were resuspended in a minimal volume of buffer A and frozen dropwise into liquid nitrogen. Cells were then lysed using a cryogenic spex mill, the lysate was allowed to thaw, and insoluble material was removed by centrifugation. The supernatant was applied to a column containing immobilized FLAG antibodies, followed by extensive washing with buffer A + 500 mM NaCl, then eluted using buffer A containing 300 mM NaCl + 2 μM FLAG peptide. To remove excess FLAG peptide, FLAG-S6-RFC was serially concentrated on a centricon 30 in buffer A containing 300 mM NaCl, then aliquoted, snap frozen in liquid nitrogen and stored at -80 °C.

RFC containing a truncated Rfc1 that lacks the N-terminal residues 1-282 ($RFC^{\Delta BRCT}$) and contains an N-terminal His/kinase tag, were co-expressed as described [3], and RFC containing the truncated Rfc1 (tRFC) was purified [1]. Briefly, *E. coli* BLR(DE3) cells with tRFC expression plasmids were grown under antibiotic selection at 30 °C to an $OD_{600}$ of ~0.6, brought to 15°C by swirling in ice water, then 1 mM IPTG was added and cells allowed to incubate a further 18 h at 15 °C. Cells were collected by centrifugation, resuspended in an equal volume of 50 mM Tris-HCl pH 7.5, 0.5 % EDTA, 5 mM DTT, 10% sucrose, 500 mM NaCl, then lysed by French press. Insoluble material was discarded by centrifugation at 4 °C. The supernatant was diluted using buffer A to bring conductivity equal to 150 mM NaCl, then applied to SP-Sepharose, washed extensively with buffer A + 150 mM NaCl and then eluted with a 300 mL gradient of buffer A + 150 mM NaCl and buffer A + 600 mM NaCl. Peak fractions eluted at ~365 mM NaCl and were pooled, diluted with Buffer A to a conductivity of 150 mM NaCl and were applied to Q-Sepharose pre-equilibrated with Buffer A + 150 mM NaCl, then eluted with a gradient of Buffer A + 150 mM NaCl to 600 mM NaCl. Peak fractions eluted at ~300 mM NaCl and were pooled, aliquoted, and stored in -80 °C.

***Polδ***

*S. cerevisiae* Polδ was expressed and purified as described [4]. Briefly, two expression vectors containing yeast pol3, pol31, and GST-pol32 plasmids were co-transformed into *E. coli* BL21(DE3) cells. 12 L of cells (1 L per 2L shaker flask) were grown at 37°C to $OD_{600}$ ~0.7 in the presence of ampicillin, kanamycin, and streptomycin. Flasks were then removed and the temperature of cultures were rapidly reduced to 15$^{o}$C by swirling on ice (monitoring temperature with a thermometer) and induced upon adding 1 mM IPTG, the incubated with shaking for a further 8 hr at 15 °C. Cells were harvested by centrifugation, resuspended in a minimal volume of 50 mM Tris-HCl pH 7.5, 1 mM EDTA, 10% glycerol, 500 mM NaCl, 2 mM PMSF. Resuspended cells were lysed in a French Press, then 30 mM spermidine was added, and the lysate was clarified by centrifugation at 23,700 × g for 1 h at 4 °C. The clarified lysate was applied to glutathione-Sepharose 4B (GE Healthcare) pre-equilibrated with 50 mM Tris-HCl pH 7.5, 10% sucrose, 500 mM NaCl. After extensive washing with the same buffer, the protein was eluted with 40 mM reduced l-glutathione, and peak fractions were pooled, mixed with 40 units of PreScission Protease to remove the GST tag from the pol32 subunit, and dialyzed overnight against 4 L of Buffer A + 300 mM NaCl. Pooled fractions were diluted with Buffer A to a conductivity equal to 80 mM NaCl then applied to a Mono-Q column equilibrated with Buffer A + 80 mM NaCl. The column was washed with 10 column volumes of Buffer A + 80 mM NaCl and was eluted with gradient of Buffer A + 80 mM NaCl to Buffer A + 200 mM NaCl. Peak fractions of the stoichiometric Polδ heterotrimer were pooled, concentrated, aliquoted, and stored in -80 °C.

***Rad27/Fen1 flap endonuclease***

S.c. Rad27 (Fen1) with a 6xHis tag at the C-terminus was cloned into pET24 (deposited to Addgene # 239204). The ScRad27-pET24 plasmid was transformed into *E. coli* BL21(DE3) cells. Cells were grown in 3L of LB with 50 μg/ml of kanamycin and grown at 37$^{0}$C to $OD_{600}$ = 0.6, then induced with 1 mM IPTG for 3 h at 37 °C. Induced cells were harvested at 5,000 X g for 15 min at 4 °C, then resuspended in 100 ml 20 mM Tris-HCl pH 7.5, 200 mM NaCl, 10% glycerol. The resuspended cells were then lysed with a French Press and cell debris was removed by centrifugation at 48,000 X g for 40 min. Next, 5 mM imidazole was added to the clarified supernatant and then the supernatant was loaded onto a 1 ml HisTrapTM HP column equilibrated in 20 mM Tris-HCl pH 7.5, 200 mM NaCl, 10% glycerol, 5 mM imidazole. The column was washed with 3 column volumes of the same buffer, then washed in steps containing either 20, 60, 100, or 150 mM imidazole in 20 mM Tris-HCl pH 7.5, 200 mM NaCl, and 10% glycerol. S.c. Rad27 (Fen1) was eluted with 6 mL 20 mM Tris-HCl pH 7.5, 200 mM NaCl, 10% glycerol containing 200 mM imidazole, diluted with 2 volumes of 20 mM Tris-HCl pH 7.5, 10% glycerol and loaded onto a 1 mL HiTrapTM Heparin HP column. Rad27 (Fen1) was eluted using a 100 to 600 mM NaCl gradient in 20 mM mM Tris-HCl pH 7.5,10% glycerol. Fractions containing the bulk of Fen1 were pooled, concentrated to 8 mg/ml, flash frozen in liquid nitrogen and stored at -80 °C.

#### *RPA*

Yeast RPA was expressed and purified as previously described [5].

#### *Fluorescent labeling*

To obtain site-specific fluorescently labeled S6-PCNA and S6-RFC, SFP synthase (4'-phosphopantetheinyl transferase) was utilized to transfer the CoA-functionalized moieties to the single serine residue contained in the tag. SFP synthase was expressed and purified as previously described [6]. S6-PCNA or S6-RFC, SFP synthase, and LD655-CoA (Lumidyne Technologies) or Cy3-CoA were incubated at a 1:2:5 molar ratio for 1 h at room temperature in the presence of 10 mM magnesium chloride. Excess dye and SFP were removed by centrifugation through a 50 kDa molecular weight cutoff Amicon Ultra Centrifugal Filter (Millipore) using Buffer A containing 300 mM sodium chloride. The resulting labeled protein was aliquoted and stored in -80 °C.

To obtain fluorescently labeled $RFC^{\Delta BRCT}$, LD555-NHS ester (Lumidyne Technologies) was used to nonspecifically label the primary amines of $RFC^{\Delta BRCT}$. Preferential N-terminal labeling was performed by labeling at low pH = 7.0 [7]. $RFC^{\Delta BRCT}$ was dialyzed out of a Tris-based buffer and into Buffer B (50 mM HEPES pH 7.0, 300 mM NaCl, 1 mM DTT, and 0.25 mM EDTA). The protein was then incubated with LD555-NHS ester at a 1:5 molar ratio for 1 h at room temperature followed by overnight at 4 °C. The reaction was quenched with 25 mM Tris-HCl pH 6.8 for 5 min. Excess dye was removed by centrifugation through a 50 kDa molecular weight cutoff Amicon Ultra Centrifugal Filter (Millipore) using Buffer B, and the labeled protein was aliquoted and stored in -80 °C.

Fluorescently labeled AF488-RPA was obtained as described previously [6]. Labeling efficiencies for all constructs ranged between 70-90%.

#### **DNA substrate preparation**

PCR of plasmid pLW58 was performed using primers PriF and PriR to amplify a 3kb AT-rich fragment and introduce two distal Nb.BbvCI nicking sites as well as cloning restriction sites. This was cloned into the 15.9-kb plasmid pRGEB32 (Addgene plasmid # 63142 [8]) via NheI/BsrGI (3-kb PCR product) and NheI/BsiWI (pRGEB32) to create the 18.9-kb plasmid pJF3KB1.

Purified pJF3KB1 was linearized by restriction digest with BsaI-HF v2 (NEB) and the resulting 20bp fragment was removed by PEG precipitation. Biotinylated oligo pairs (Oli1+Oli2, Oli3+Oli4) were annealed and ligated (T4 DNA ligase, NEB) to the two BsaI overhangs of the linearized plasmid. Ligase was then heat inactivated. The DNA was nicked on the same strand at the two nicking sites at either end of the 3kb AT-rich region (Nb.BbvCI, NEB). The endonuclease was then heat inactivated and the DNA was purified away from excess oligos by PEG precipitation. Aliquots of the intermediate linearized pJF3KB1 and the final biotinylated/nicked substrate were digested with SphI and SpeI (NEB) to check for successful BsaI restriction, PEG purification, and introduction of the oligo pairs at both ends of the DNA. The substrate was stored in TE buffer at 4 °C. All synthetic oligonucleotides were ordered from Integrated DNA Technologies:

PriF: AAAAA*GCTAG*CCTCAGCATATTAGCTAAAACTAAAAGTGGTAAAACG (italic = NheI, underline = Nb.BbvCI)
PriR: AAAAA*TGTACA*GCTGAGGCGGAATCGGTAGTAAGTTATAGC (italic = BsrGI, underline = Nb.BbvCI)
Oli1: 5’ dual-biotin-CAAGCCGCCATTCCACTCTGCCTA
Oli2: 5’ dual-biotin-CAAGCCGCCATTCCACTCTGCCTA
Oli3: 5’ dual-biotin-CTTGCTCGTAGTCAATGCGTCAC
Oli4: 5’ phosphate-GGCAGTGACGCATTGACTACGAGCAAG

**Single-molecule experiments**

***Experimental setup***

Single-molecule experiments were performed on a LUMICKS C-Trap instrument at room temperature. This instrument combines three-color confocal fluorescence microscopy with dual-trap optical tweezers [9]. Data was acquired using LUMICKS Bluelake software version 1.6.16. Rapid optical trap movement was achieved using a computer-controlled stage in a flow cell containing 5 separate channels. Channels 1-3 were separated by laminar flow and were used to tether DNA between two 4-μm streptavidin-coated polystyrene beads (Spherotech) held in optical traps. In channel 1, a single bead was caught in each trap. The traps were next moved to channel 2, and biotinylated DNA was tethered between the two beads as denoted by an increase in the force reading. The tether was then moved into channel 3, and the flow was stopped. The presence of a single DNA tether was confirmed by the generated force-distance curve.

To generate a gapped DNA tether, the ds gapped DNA construct was pulled to high forces (~60 pN) in a no-salt buffer (Tris-HCl pH 8.0). Flow was turned on (~0.1-0.2 bar) for 5 s and the tether was relaxed (**Figure S1B**). Flow was turned off, and successful generation of the gapped tether was confirmed by the force-distance curve (**Figure S1C**).

Channels 4 and 5 were loaded with proteins as described for each assay. Flow was turned off when collecting data and visualizing protein behavior. For fill-in assays, either the force or distance between the beads was held constant, and the resultant change in either the trap position or the force was recorded as a measurement of fill-in rate and completion.

***Fluorescence detection***

Cy3, LD655, and AF488 fluorophores were excited by three laser lines at 532, 638, and 488 nm respectively. Confocal line scanning at 100 ms/line through the center of each bead was used to generate kymographs. Occasionally, individual lasers were turned off to confirm the presence of fluorescently labeled proteins.

To investigate the behavior of LD655-PCNA and Cy3-RFC or LD555-RFC$^{\Delta BRCT}$, optical traps tethering a gapped or λ DNA molecule (LUMICKS) under 6 or 1 pN of constant tension respectively were moved into channel 4 of the microfluidic flow cell containing 8 nM of each protein plus 100 nM of RPA in Image Buffer (25 nM Tris-HCl pH 7.5, 100 nM sodium chloride, 10 mM magnesium acetate, 3 mM DTT, 2 mM TCEP, 1% glycerol, 40 μg/uL BSA, 0.1 mM EDTA, 4 mM ATP, 1 mM cyclooctatetraene [Sigma], 1 mM 4-nitrobenzyl alcohol [Sigma], and oxygen scavenging reagents 10 nM protocatechuate-3,4-dioxygenase [OYC Americas] and 2.5 mM protocatechuic acid

[Sigma]). For high salt challenge experiments, the tether with bound proteins was moved to channel 5 with Image Buffer containing 500 mM NaCl.

For fill-in experiments, a gapped DNA tether was moved to channel 4, which contained 20 nM Polδ, 5 nM LD655-PCNA, 5 nM Cy3-RFC or LD555-RFC$^{\Delta BRCT}$, and 100 nM RPA in Image Buffer supplemented with 115 μM of all deoxynucleotides and held at the specified tension. When RPA was visualized, 20 nM of AF488-RPA was used along with 5 nM of dark PCNA and RFC in lieu of the fluorescently labeled constructs. For experiments with FEN1, 15 nM of FEN1 was added to each sample.

#### *Data analysis*

Raw position data of righthand bead (Trap 1) collected at 100 Hz was first smoothed with a Savitzky-Golay smoothing filter of polynomial order 3 and frame length of 501 data points [10]. To infer polymerase trajectory from bead position data, the DNA fill-in reaction was modeled using a linear combination of the freely-jointed chain model for the single-stranded segment and the extensible worm-like chain model for the double-stranded segment [11, 12]. From this linear combination, we predict the change in tether length per base pair filled in as a function of force applied. This relation was used to convert the bead trajectory in nanometers to a fill-in trajectory in base pairs. The instantaneous velocity was extracted from the fill-in trajectory with the gradient function, which returns the one-dimensional numerical gradient of the input vector. Pauses in the trajectory were identified using an empirical velocity cutoff of +15 bp/s. To avoid over-counting from local rapid changes in velocity, we only identify pauses at least 1 second long, and if pauses are separated by intervals of less than 2 seconds, the pauses are joined together. All analyses were performed in MATLAB R2024b.

#### Structural prediction

To structurally predict the PCNA-RFC-Polδ complex, the AlphaFold webserver featuring AlphaFold3 was used (alphafoldserver.com). Three copies of *S. cerevisiae* PCNA protein sequence (POL30) (*Saccharomyces* Genome Database [SGD] #: YBR088C), one copy of RFC1-5 (SGD #: YOR217W; YJR068W; YNL290W; YOL094C; YBR087W), and one copy of each of the Polδ subunits, POL3 (SDG #: YDL102W), POL31 (SDG #: YJR006W), and POL32 (SDG #: YJR043C) were inputted as well as the two complementary single-stranded DNA sequences (Primer strand 5' to 3': AGCTATGACCATGATTACGAATTGC; Template strand 5' to 3': CTGCACGAATTAAGCAATTCGTAATCATGGTCATAGCT) [13]. The highest rank model was chosen for visualization and atoms were hidden if they had a B-factor less than 0.65.

#### Mass photometry

Data were collected using a OneMP mass photometer (Refeyn). Calibration of the instrument was performed using bovine serum albumin (66 kDa), beta amylase (224 kDa), and thyroglobulin (670 kDa). Movies were acquired for 6,000 frames (60 s) using AcquireMP software (version 2.4.0) under the default settings. Protein concentrations were empirically established to achieve ~75 binding events per second. Peak mass values are predicted to exist within ~5% error. Raw data were converted to frequency distributions using Prism 9 (GraphPad) using a bin size of 10 Da.

**Genetic methods**

***Yeast strains***

Strains used in this study are listed in **Table S1** and are isogenic to a *RAD5* derivative of W303 (*MATa ade2-1 can1-100 ura3-1 his3-11,15, leu2-3, 112 trp1-1 rad5-535*) [14]. At least two strains per genotype were examined for each assay and only one is listed in **Table S1**. Mutant construction and protein tagging were conducted using standard PCR-based method and genetic engineered loci were verified by sequencing. Standard procedures were used for cell growth, media preparation, genetic crosses, tetrad analyses, and genotoxin treatment. Cells were grown at 30 °C in YPD media for both tetrad analyses and genotoxin treatment experiments.

Primers used in the yeast genetics experiments:

Rfc1 1789 F: 5'-GCTTGTTAAGAAAGAAGAGG

Rfc1 1753 F: 5'-GGCTAATTGGGAAAATTCAAAG

1315 F: 5'-GGCTCAAGATGTACTAGATA

1290 F: 5'-GTTCAGACTCGGCTTCGAAT

Rfc1 Far DR: 5'-GTTCAGCAAAGGCATCCCTT

Rfc1 DR: 5'-CTGTACGTACAACCGTCGTT

***Protein level examination***

To detect the wild-type and mutant Rfc1 proteins C-terminal tagged with the TAP tag, protein extracts were made using a TCA (Trichloroacetic acid) method as previously described [15]. In brief, cells pellets were resuspended in 20% TCA and homogenized using glass beads in a FastPrep-24 bead beating instrument (MP Biomedicals). The lysate was centrifuged to remove supernatant. The precipitated proteins were dissolved in Laemmli buffer (65 mM Tris-HCl pH 6.8, 2% SDS, 10% glycerol, 5% β-mercaptoethanol, and 0.025% bromophenol blue) with 2 M Tris to neutralize the lysate. Prior to loading, samples were boiled for 5 min and spun down to remove insoluble materials. Samples were separated on NuPAGE™ 4–12% Bis-Tris gels (Thermo Fisher NP0322) for immunoblotting to detect the TAP-tagged Rfc1 proteins.

**Statistical analysis**

Errors reported in this study represent standard deviation (SD) or standard error of the mean (SEM). *P* values were determined from two-tailed unpaired *t* tests (with Welch's correction as specified in each figure caption) for comparison between two conditions using Prism 10 (GraphPad).

**Data availability**

Source data (raw values, uncropped gels) are provided with this paper. All kymographs used for analysis have been deposited as datasets in Zenodo.

**Code availability**

Kymographs were processed and analyzed using a custom script that can be accessed on LUMICKS Harbor (https://harbor.lumicks.com/single-script/c5b103a4-0804-4b06-95d3-20a08d65768f).

### SUPPLEMENTARY FIGURES

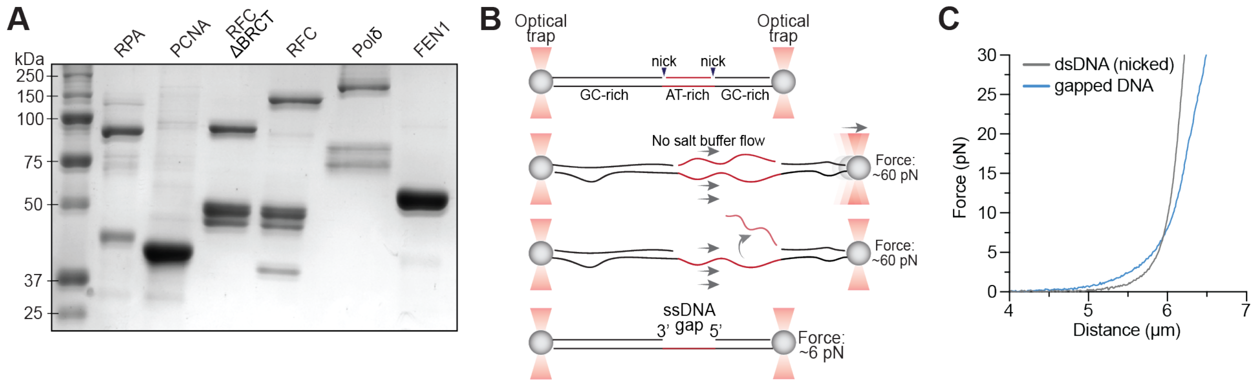


**Figure S1. Protein and DNA substrate preparations, related to Figures 1 and 3.**
(**A**) SDS-PAGE gel showing each purified protein used in this work.
(**B**) Schematic of the protocol to generate a gapped DNA tether in situ in an optical tweezers setup. A single dsDNA molecule containing a 3-kb, AT-rich segment flanked by two nicks and a 10-kb and 6-kb GC-rich arm was pulled to high forces (~60 pN). A no-salt buffer was flowed in gently to remove the untethered AT-rich ssDNA strand. The tether was then relaxed, and the flow was turned off, leaving a 3-kb ssDNA gapped DNA tether for subsequent single-molecule imaging.
(**C**) Force-distance curves for dual-nicked dsDNA (grey) and 3-kb gapped DNA (blue) tethers. The differences allow the assessment of successful gap generation.

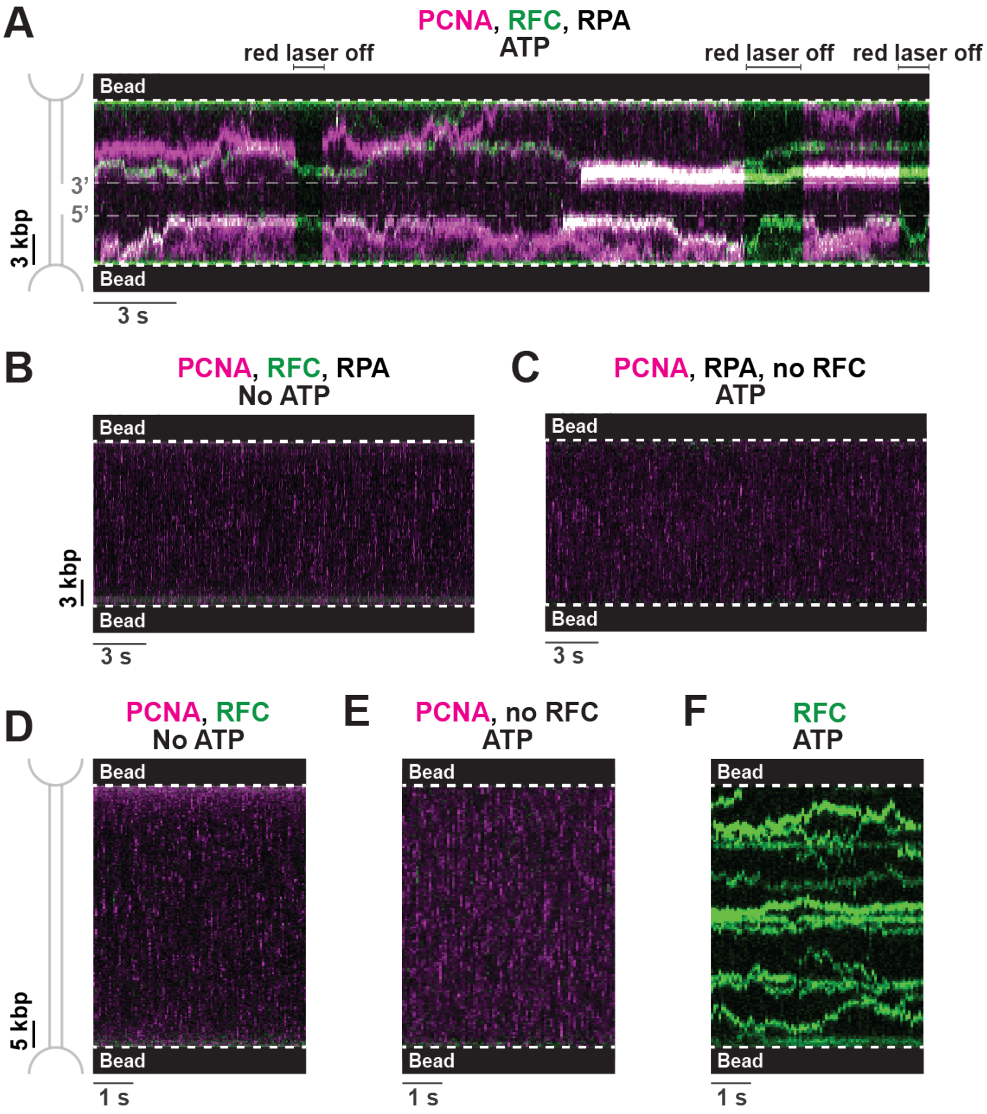


**Figure S2. Additional experiments showing PCNA and RFC behavior on gapped or duplex DNA, related to Figures 1 and 2.**

(**A**) Representative kymograph showing a gapped DNA tether incubated with 8 nM LD655-PCNA, 8 nM Cy3-RFC, and 100 nM RPA with ATP. Red laser was turned off briefly to confirm the presence of green RFC signals.

(**B**) Representative kymograph showing a gapped DNA tether incubated with 8 nM LD655-PCNA, 8 nM Cy3-RFC, and 100 nM RPA in the absence of ATP.

(**C**) Representative kymograph showing a gapped DNA tether incubated with 8 nM LD655-PCNA and 100 nM RPA with ATP in the absence of RFC.

(**D**) Representative kymograph showing a ds λ DNA tether incubated with 8 nM LD655-PCNA, 8 nM Cy3-RFC, and 100 nM RPA in the absence of ATP.

(**E**) Representative kymograph showing a ds λ DNA tether incubated with 8 nM LD655-PCNA and 100 nM RPA with ATP in the absence of RFC.

(**F**) Representative kymograph showing a ds λ DNA tether incubated with 8 nM Cy3-RFC (without PCNA) and 100 nM RPA with ATP.

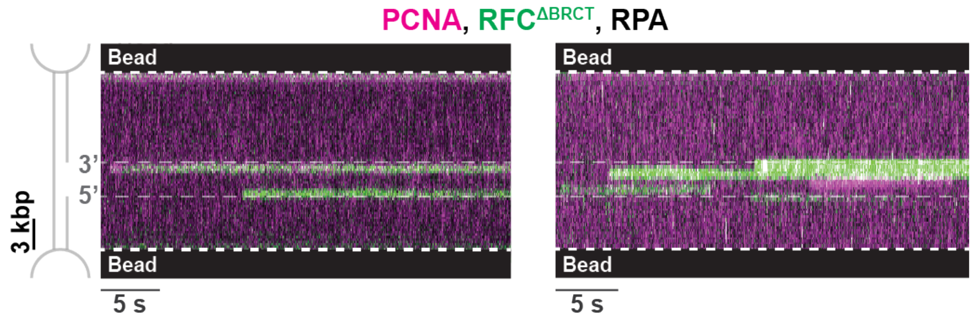


**Figure S3. RFC$^{\Delta BRCT}$ binds to RPA-coated ssDNA, related to Figure 3.**
Two representative kymographs showing gapped DNA tethers incubated with 8 nM LD655-PCNA, 8 nM LD555-RFC$^{\Delta BRCT}$, and 100 nM RPA with ATP.

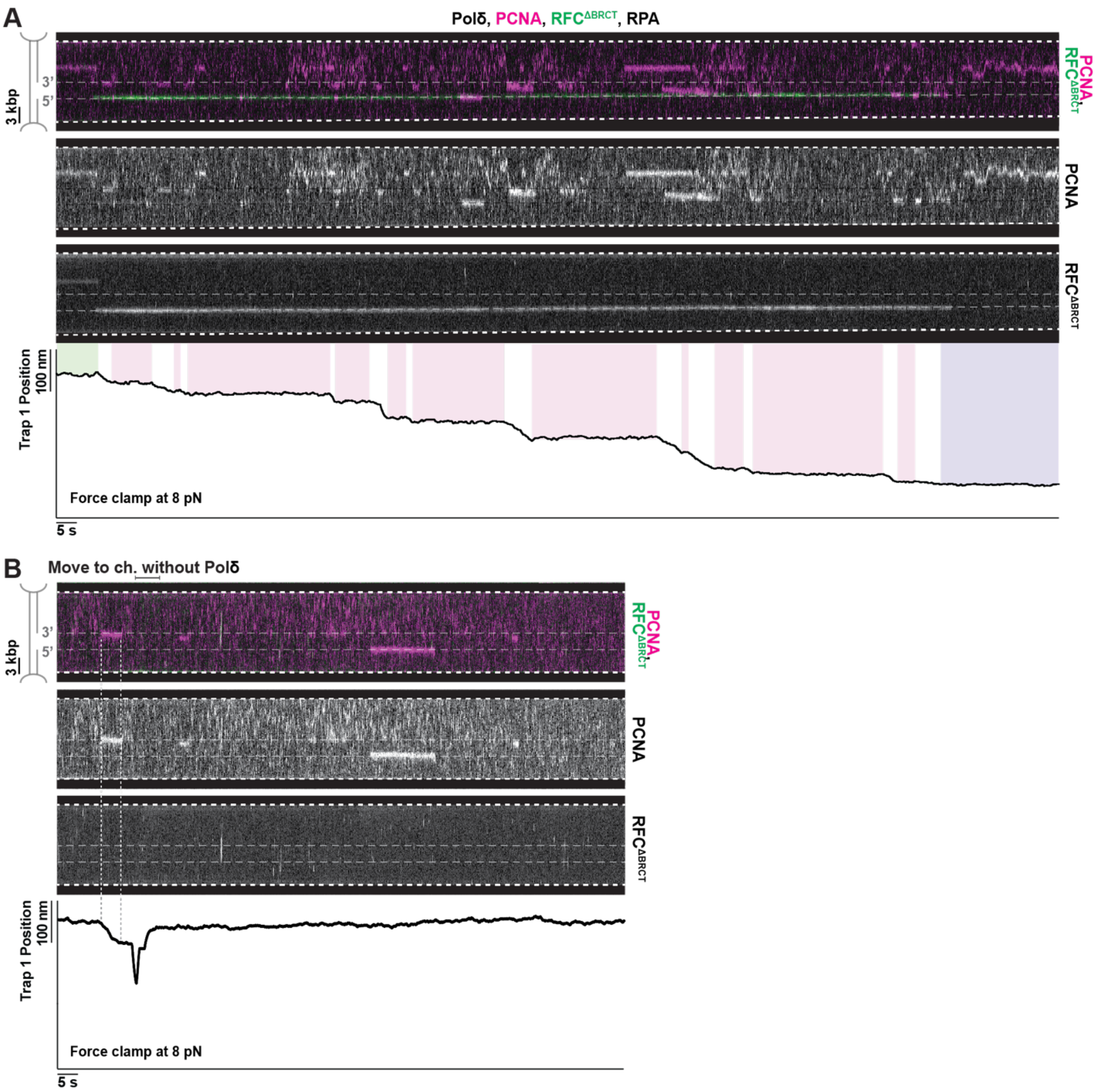


**Figure S4. Additional data showing PCNA and RFC$^{\Delta BRCT}$ behaviors during fill-in synthesis, related to Figure 4.**

(**A**) Kymograph and associated trap position changes at a constant force of 8 pN showing fill-in reaction of a gapped DNA tether incubated with 20 nM Polδ, 5 nM LD655-PCNA, 5 nM LD555-RFC$^{\Delta BRCT}$, and 100 nM RPA. The PCNA and RFC channels are shown in separate kymographs to confirm the presence of each signal. Pink boxes in the bottom row denote periods of inactive synthesis, green box denotes the period prior to the start of fill-in, and purple box denotes the completion of fill-in.

(**B**) Kymograph and associated trap position changes at a constant force of 8 pN where the fill-in reaction initiated on the gapped DNA tether (denoted by the vertical dotted lines) in a channel containing the same components as (**A**). The partially filled-in tether was then moved to a different channel containing the same components except Polδ. The fill-in reaction failed to resume, indicating the requirement of free Polδ in solution for continued synthesis.

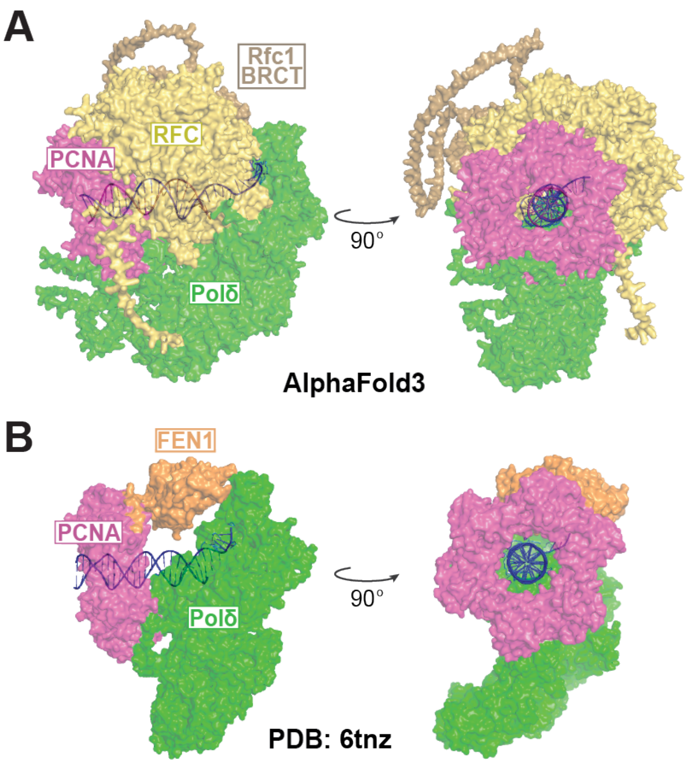


**Figure S5. Structural models of PCNA-Polδ assemblies with an additional PCNA-binding protein, related to Figures 4 and 5.**

(**A**) AlphaFold3 prediction of a complex containing PCNA, RFC, Polδ, and a 3' recessed DNA.

(**B**) Cryo-EM structure of the PCNA-FEN1-Polδ ternary complex with a 3' recessed DNA (PDB: 6tnz) [16].

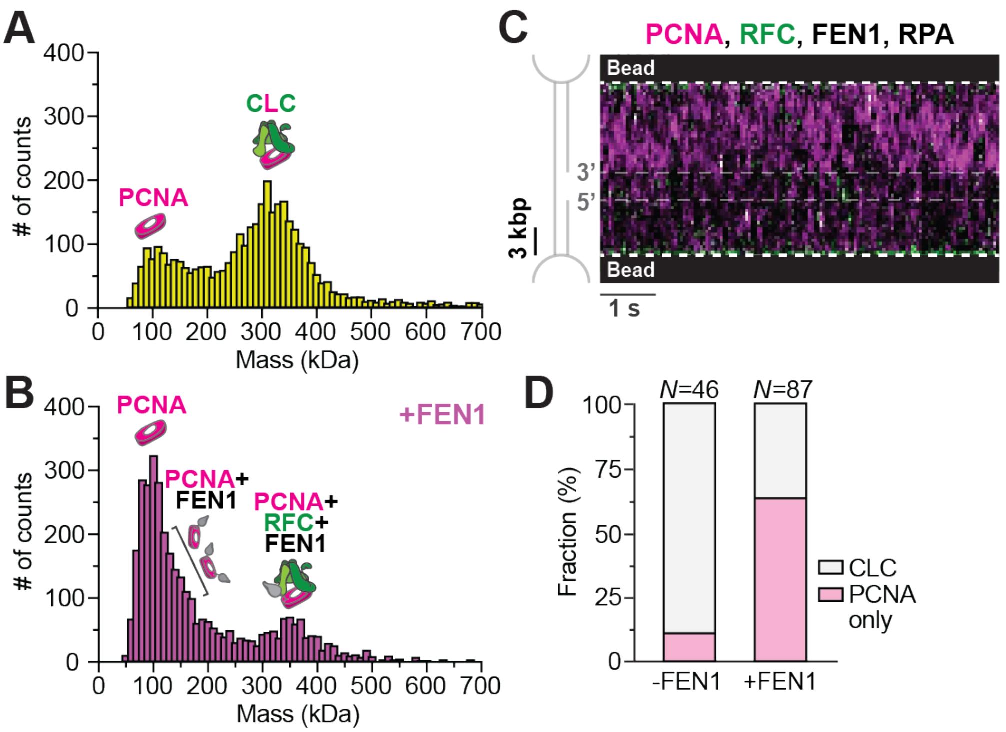


**Figure S6. FEN1 and RFC compete for PCNA binding, related to Figure 5.**

(**A**) Mass distribution for PCNA and RFC incubated in solution with ATPγS. Predicted complexes for the mass peaks are indicated.

(**B**) Mass distribution for PCNA and RFC incubated in solution with ATPγS in the presence of excess FEN1.

(**C**) Representative kymograph showing a gapped DNA tether incubated with 8 nM LD655-PCNA, 8 nM Cy3-RFC, 15 nM FEN1, and 100 nM RPA with ATP.

(**D**) Fraction of trajectories on gapped DNA corresponding to CLC (colocalized PCNA and RFC signals) versus PCNA only (no RFC signal) in the absence ($N$ = 46 from 6 independent tethers) or presence ($N$ = 87 from 11 independent tethers) of FEN1.

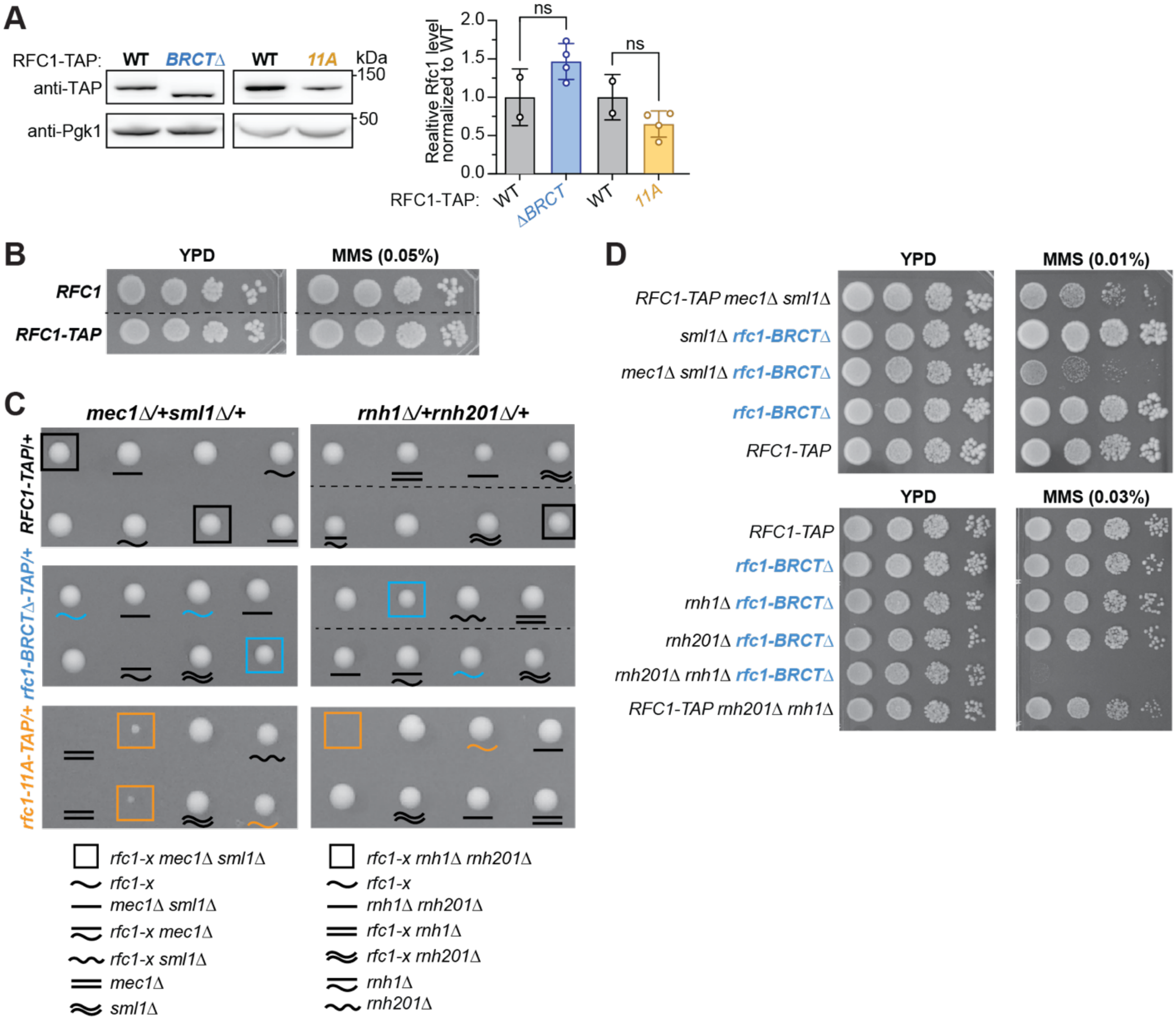


**Figure S7. Genetic analyses of Rfc1 BRCT mutants, related to Figure 6.**

(**A**) Immunoblotting (left) and quantification (right) of protein levels for Rfc1-BRCTΔ (deleted residues: 154-230) and Rfc1-11A (mutated residues: T166A, R174A, R187A, T189A, K190A, S191A, S193A, S194A, K195A, K208A, K209A) compared to their wild-type counterpart. All examined proteins were C-terminally tagged with a TAP tag. Pgk1 was used as a loading control. Quantification was derived from at least two spore clones per genotype. The relative protein levels of Rfc1 over Pgk1 were calculated and normalized to the values for the wild-type. Bars represent mean and SD. Significance was calculated using a two-tailed unpaired t-test.

(**B**) Wild-type Rfc1 tagged with TAP did not lead to genotoxic sensitivity of *S. cerevisiae* cells. The dotted line separates cells located in different sections of the same spot assay plate.

(**C**) Genetic interactions of *rfc1-BRCTΔ and rfc1-11A* with mutants lacking the Mec1 protein or the RNase H1 and RNase H2 proteins using tetrad analyses. The dotted line separates tetrads located in different sections of the same dissection plate.

(**D**) Genetic interactions of *rfc1-BRCT*Δ with Mec1 or RNase H1 and RNase H2 mutants in the absence or presence of MMS.

**Table S1. Yeast strains, related to Figure 6.**

All strains are isogenic to a *RAD5* derivative of W303 (*MATa ade2-1 can1-100 ura3-1 his3-11,15, leu2-3, 112 trp1-1 rad5-535*). Only one strain is listed per each genotype and at least two independent isolates of each genotype were used in the experiments.

| Strain | Genotype | Source | Figures |
|---|---|---|---|
| G16 | *MATa ade2-1 can1-100 ura3-1 his3-11,15, leu2-3, 112 trp1-1 rad5-535* | Lab collection | S7B |
| T799-1 | *MATα RFC1-TAP::HIS* | Lab collection | 6A, S7A, S7B, S7D |
| T2338-2-1B | *MATa rfc1-BRCTΔ-TAP::HIS* | This study | 6A, S7A, S7D |
| X9587-1D | *MATa rfc1-11A-TAP::HIS* | This study | 6A, S7A |
| X9673-5B | *MATα rnh1Δ::hphNT1 rnh201Δ::KAN RFC1-TAP::HIS* | This study | S7D |
| X9682-1C | *MATa rnh1Δ::hphNT1 rnh201Δ::KAN rfc1-BRCTΔ-TAP::HIS* | This study | S7D |
| X9682-6A | *MATα rnh201Δ::KAN rfc1-BRCTΔ-TAP::HIS* | This study | S7D |
| X9682-8B | *MATa rnh1Δ::hphNT1 rfc1-BRCTΔ-TAP::HIS* | This study | S7D |
| X9671-1D | *MATa mec1Δ::TRP sml1Δ::HIS RFC1-TAP::HIS* | This study | S7D |
| X9680-1D | *MATa mec1Δ::TRP sml1Δ::HIS rfc1-BRCTΔ-TAP::HIS* | This study | S7D |
| X9680-4A | *MATα sml1Δ::HIS rfc1-BRCTΔ-TAP::HIS* | This study | S7D |
| X9724 | rad27Δ::URA3 *rfc1-BRCTΔ-TAP::HIS* | This study | 6B |
| X9755 | rad27Δ::URA3 *RFC1-TAP::HIS* | This study | 6B |
| X9723 | rad27Δ::URA3 *rfc1-11A-TAP::HIS* | This study | 6B |
| X9752 | *rad51Δ::URA3 RFC1-TAP::HIS* | This study | 6C |
| X9685 | *rad51Δ::URA3 rfc1-BRCTΔ-TAP::HIS* | This study | 6C |
| X9684 | rad51Δ::URA3 *rfc1-11A-TAP::HIS* | This study | 6C |
| X9671 | *mec1Δ::TRP sml1Δ::HIS RFC1-TAP::HIS* | This study | S7C |
| X9680 | *mec1Δ::TRP sml1Δ::HIS rfc1-BRCTΔ-TAP::HIS* | This study | S7C |
| X9590 | *mec1Δ::TRP sml1Δ::HIS rfc1-11A-TAP::HIS* | This study | S7C |
| X9673 | *rnh1Δ::hphNT1 rnh201Δ::KAN RFC1-TAP::HIS* | This study | S7C |
| X9682 | *rnh1Δ::hphNT1 rnh201Δ::KAN rfc1-BRCTΔ-TAP::HIS* | This study | S7C |
| X9589 | *rnh1Δ::hphNT1 rnh201Δ::KAN rfc1-11A-TAP::HIS* | This study | S7C |

### SUPPLEMENTARY REFERENCES


1. Zheng, F., et al., *Cryo-EM structures reveal that RFC recognizes both the 3'- and 5'-DNA ends to load PCNA onto gaps for DNA repair.* Elife, 2022. **11**.
2. Georgescu, R.E., et al., *Mechanism of asymmetric polymerase assembly at the eukaryotic replication fork.* Nat Struct Mol Biol, 2014. **21**(8): p. 664-70.
3. Finkelstein, J., et al., *Overproduction and analysis of eukaryotic multiprotein complexes in Escherichia coli using a dual-vector strategy.* Analytical Biochemistry, 2003. **319**(1): p. 78-87.
4. Langston, L.D. and M. O'Donnell, *DNA polymerase delta is highly processive with proliferating cell nuclear antigen and undergoes collision release upon completing DNA.* J Biol Chem, 2008. **283**(43): p. 29522-31.
5. Henricksen, L.A., C.B. Umbricht, and M.S. Wold, *Recombinant replication protein A: expression, complex formation, and functional characterization.* J Biol Chem, 1994. **269**(15): p. 11121-32.
6. Wasserman, M.R., et al., *Replication Fork Activation Is Enabled by a Single-Stranded DNA Gate in CMG Helicase.* Cell, 2019. **178**(3): p. 600-611.e16.
7. Sélo, I., et al., *Preferential labeling of alpha-amino N-terminal groups in peptides by biotin: application to the detection of specific anti-peptide antibodies by enzyme immunoassays.* J Immunol Methods, 1996. **199**(2): p. 127-38.
8. Xie, K., B. Minkenberg, and Y. Yang, *Boosting CRISPR/Cas9 multiplex editing capability with the endogenous tRNA-processing system.* Proc Natl Acad Sci U S A, 2015. **112**(11): p. 3570-5.
9. Candelli, A., G.J. Wuite, and E.J. Peterman, *Combining optical trapping, fluorescence microscopy and micro-fluidics for single molecule studies of DNA–protein interactions.* Physical Chemistry Chemical Physics, 2011. **13**(16): p. 7263-7272.
10. Savitzky, A. and M.J.E. Golay, *Smoothing and Differentiation of Data by Simplified Least Squares Procedures.* Analytical Chemistry, 1964. **36**(8): p. 1627-1639.
11. Smith, S.B., Y. Cui, and C. Bustamante, *Overstretching B-DNA: the elastic response of individual double-stranded and single-stranded DNA molecules.* Science, 1996. **271**(5250): p. 795-9.
12. Odijk, T., *Stiff Chains and Filaments under Tension.* Macromolecules, 1995. **28**(20): p. 7016-7018.
13. Zheng, F., et al., *Structure of eukaryotic DNA polymerase δ bound to the PCNA clamp while encircling DNA.* Proc Natl Acad Sci U S A, 2020. **117**(48): p. 30344-30353.
14. Zhao, X. and G. Blobel, *A SUMO ligase is part of a nuclear multiprotein complex that affects DNA repair and chromosomal organization.* Proc Natl Acad Sci U S A, 2005. **102**(13): p. 4777-4782.
15. Wan, B., et al., *Mms22-Rtt107 axis attenuates the DNA damage checkpoint and the stability of the Rad9 checkpoint mediator.* Nat Commun, 2025. **16**(1): p. 311.
16. Lancey, C., et al., *Structure of the processive human Pol δ holoenzyme.* Nature Communications, 2020. **11**(1): p. 1109.
